# An Enzymatic Active Site Embedded in a DNA Nanostructure

**DOI:** 10.1101/804518

**Authors:** Tadija Kekic, Yasaman Ahmadi, Ivan Barisic

## Abstract

Artificial enzymes hold a great potential in the field of biotechnology. The currently available design strategies are limited to recruit and organise secondary and tertiary structural elements, and functional motifs of native enzymes. Here, we demonstrate an approach towards the bottom-up development of an artificial enzyme based on DNA nanotechnology. Structural analyses of the FlaI ATPase were used to identify a set of amino acids essential for the catalytic reaction. The selected amino acids were used to design four peptide-oligonucleotide conjugates (POCs). The POCs were integrated into a DNA nanostructure and reconstituted the catalytic site of FlaI. DNA origami technology was employed to maintain the relative orientations of the amino acids and their positions analogous to those in the protein catalytic site. The substrate turnover rates were found to be comparable in our artificial nanozyme (k_cat_ = 0.3622 s^-1^) and the native enzyme (k_cat_ ≈ 0.016 s^-1^). The nanozymes could be recycled using an ultrafiltration protocol and reused in multiple experiments allowing a reliable reproduction of the measurements. The emulation of enzymatic activity, by this novel and computable technological framework, can be of high relevance in many different areas.

---

Biological systems employ enzymes to catalyse thousands of chemical reactions with a speed and efficiency rarely achievable by artificial processes. The use of enzymes, however, is often limited by factors such as the appropriate expression system, complexation with partners, and correct phosphorylation and glycosylation profiles [1-3]. The isolation of enzymes is often challenging, and low yields and poor purity lead to high costs. To avoid protein isolation difficulties, evolutionary conserved structural motifs of enzymes can be used to design artificial peptides [18]. In comparison to the native enzymes, the speed of peptide’s fabrication is significantly higher. However, studies have reported low catalytic activity, aggregation problems, and the inability to recreate the tertiary structure of the active site.

The need for enzymes for catalysis of various chemical and biological reactions has inspired the development of biomimetic materials with catalytic properties [4-15]. To achieve *de novo* catalytic properties several design strategies have been proposed [7-9, 11-14, 18-19]. The *molecular imprinting* approach is used to embed the key functional groups of the idealised catalytic site in a polymer matrix. The placement of the functional groups is predefined by their interactions with the transition state analogue (TSA), thus forming a catalytic site cavity. TSAs can also be used as targets for the development of *catalytic antibodies*. The fabrication of TSA-based, catalytically active material is very fast. However, studies have reported low efficiency in comparison to the native enzymatic catalysis [12-13].

An alternative *de novo* approach is the integration of idealised catalytic site’s residues into a well-defined protein framework. To reduce undesirable spatial flexibility of the catalytically active amino acids, the framework structures are based on proteins with highly conserved and stable tertiary structures [7]. The enzymatic activity can be further optimised experimentally by the introduction of mutations. The reported catalytic activity of these systems is similar to native enzymes and the isolation process is simplified by choosing the adequate protein scaffold. Using this approach, the direct integration of the active site into the protein scaffold limits the design process, because only a few key amino acids instead of whole secondary or tertiary motifs can be integrated in the catalytic site.

With the advancement of the DNA origami technology, DNA emerged as a versatile material for generating highly complex nanostructures of unprecedented precision [20-22]. The integration of glucose oxidase (GOx) and horseradish peroxidase (HRP) enzymes, as a reliable multi-enzyme system [17], on the surface of DNA nanostructures has been used to create enzymatic cascades with improved catalytic efficiency [5]. In addition, the encapsulation of an enzyme into a nanostructure also allows a precise regulation of the enzyme activity due to a more tunable molecular environment [10, 16]. However, this approach is limited by the properties of the enzymes, and the ability to isolate them in the active form.

In this study, we showcase a bottom-up approach for the development of *de novo* catalytic properties by emulating the catalytic site of an ATP-driven motor protein into a DNA-based nanostructure. The emulation of the ATPase activity was based primarily on the FlaI protein complex from *Sulfolobus acidocaldarius*, the simplest transmembrane motoric supercomplex [3]. The hydrolysis of ATP in FlaI, accompanied by the resulting domain movements in the protein complex, generates a torque and rotational force that is channelled through the filament of the transmembrane protein supercomplex of the archaellum [3, 23]. In addition, this FlaI was selected due to a set of criteria, such as an elucidated crystal structure, a well-defined and open active site, available reaction kinetics data, and an established colorimetric assay for tracing the substrate turnover [3, 23, 24].

## Analysis of FlaI protein complex

The crystal structure of the FlaI hexamer (PDB: 4IHQ) was used to analyse the protein complex *in silico* [3]. Domain dynamics and the interactions between the enzyme’s active site, the substrate, the products, and the cofactor were investigated using molecular dynamics (MD) simulations (Fig. 1.A-C). Due to the highly dynamic nature of the FlaI hexamer, we observed significant deviations in the conformations and relative positions of the amino acids in the active site (Fig. S10). The analysis of the MD simulations and the available quantum mechanics data from similar systems [27 - 30] helped to identify and investigate the key amino acids, which are both structurally and catalytically important. The final amino acids that were embedded in the DNA nanostructure were also selected by the comparison of phylogenetic and structural data from homologous proteins (Fig 1.D-I, and Fig. S1-S3). The key amino acids were assigned to four groups, based on their role in the interaction with the substrate, the products, and the cofactor (Fig. 2). The first group is mainly responsible for the substrate binding capabilities in the surrounding area of the nucleotide base and deoxyribose sugar. The second group is made of residues of the conserved Walker B motif and interacts with the metallic ion in the proximity of the gamma phosphate of the nucleotide. The third group resembles the environment surrounding the gamma phosphate, with direct involvement in the catalytic process. Finally, the fourth group comprises a conserved Walker A motif and LRLR nest.

**Fig. 1.**
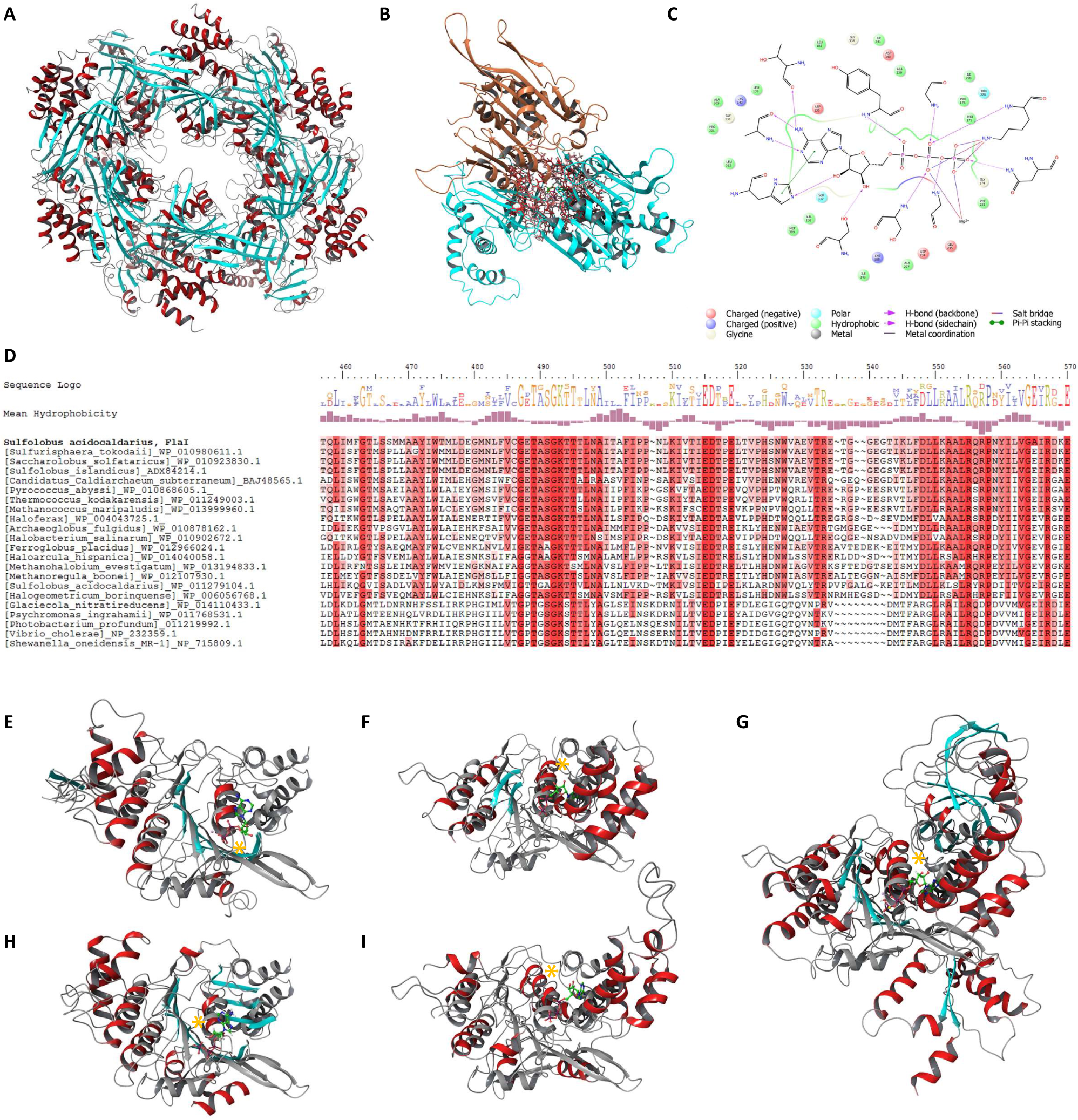
The structural phylogenetic analysis of the FlaI family and FlaI homologues with conserved catalytic motifs. The selection of the key amino acids and catalytic motifs. A) The FlaI hexamer, based on the crystal structure (PDB: 4IHQ) and obtained after 200 ns of molecular dynamics (MD) simulation. Alpha helices and B-sheets are colored in red and cyan, respectively; B) Structure of the three FlaI domains involved in a single active site. The three domains belong to two neighboring FlaI monomers colored in blue and orange, respectively; C) Two-dimensional representation of the interactions between substrate, cofactor, and amino acids in the active site of the FlaI protein. Interacting amino acids are represented with their chemical structure, while the neighboring amino acids are color-coded based on their influence on the environment; D) Segment of the protein sequence alignment of the FlaI family, alignment sequence logo and the mean hydrophobicity with the conserved motifs and key amino acids used in the modelling of the emulated catalytic site. The white-red color gradient represents the conservation level of each residue. E-I) The structures of the FlaI homologues aligned to the reference structure of the FlaI’s active site obtained from the MD simulation. Alpha helices and beta sheets are colored in red and blue respectively, while the reference structure is colored in gray. The highest alignment score, obtained in the region of Walker A motifs, is marked by the yellow asterix. The crystal structures of the FlaI homologues are available under the PDB codes: 1G6H, 2ZAN, 5C18, 4RVC, 5Z6R.

**Fig. 2.**
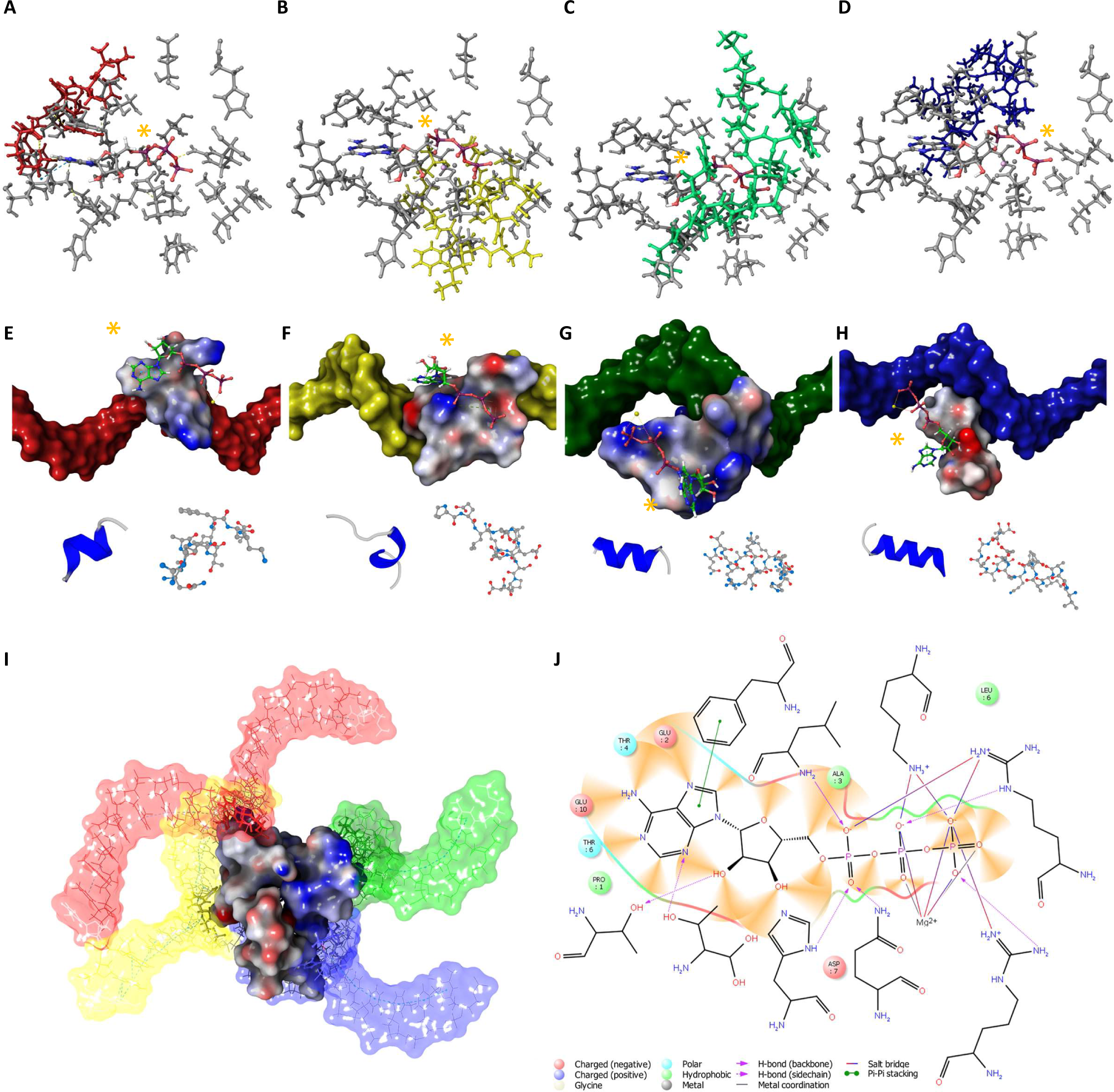
The modelling of the emulated active site of the DNA Scaffold-Embedded Protein Emulation Complex (D-SEPEC). Main steps in modelling of the emulated active site. A-D) Through the process of addition of neighboring residues, the key amino acids of the catalytic center of the FlaI enzyme were merged into four peptides, 10 to 15 amino acids in length. These peptides served as a base for the development of the peptide-oligonucleotide conjugates (POC). The peptides of POC 1 to 4 are depicted with licorice structures in designated color: red, yellow, green and blue, respectively. The key amino acids of the POC elements are aligned with the native catalytic site (see Fig. S4). The position of the substrate, ATP, is marked with the yellow asterix; E-H) The peptides of POC 1 to 4 linked with the appropriate oligonucleotide formed distinct interactions with the substrate. The van der Waals (VDW) surfaces of the peptides are colored based on partial charges, while the surfaces of oligonucleotides are in designated colors for each POC. The substrate, ATP, is represented by a licorice model. The most frequently observed conformation cluster of each peptide element, obtained by the folding simulations are shown in both atomic and cartoon representation; I) The POC elements combined to form the emulated catalytic site are represented by their VDW surfaces, colored in the partial charges. The oligonucleotide elements, represented by both transparent VDW surface and atomic representations in designated colors, retained their ds DNA conformation by hybridisation with the short Carrier Strand fragments (not shown). Following the optimisation process, this model was used for the integration into the DNA-origami nanostructure. J) Two-dimensional representation of the positioning and interactions between substrate, cofactor, and amino acids in the emulated catalytic center of the D-SEPEC nanozyme. Interacting amino acids are represented by their chemical structure, and neighboring amino acids are color-coded based on their influence on the environment.

## Design of the artificial catalytic site

The obtained structural data was used for the *in silico* modelling of a DNA Scaffold-Embedded Protein Emulation Complex or D-SEPEC. The core structure comprising the key amino acids was gradually expanded by the systematic, *in silico*, addition of neighbouring amino acids, while keeping the elements of the D-SEPEC close to the steric and electrostatic energy minimum (Fig. 2.A-D, and Fig. S4) [31]. Throughout this process, the conformation of amino acids was adjusted from the one obtained by the MD simulations of the native protein. A residue-type adjustment of amino acids was in some cases necessary to retain similar functional properties with respect to the difference in the environment and the interactions of the amino acids within the active site. Further modifications of the key amino acids were inserted to enhance the stability of the binding interactions between the substrate and the active site, or to adjust the amino acid residues involved in the hydrophobic interactions of the original protein (Table S2). In the expanded model, neighbouring amino acids were joined by peptide bonds, until four oligopeptides, 10 – 15 amino acids in length, were formed. A total of 46 amino acids from the FlaI protein were used. The D-SEPEC model was further expanded by the addition of four oligonucleotides, each having a single base modification (5-ethynyl-2’deoxyuridine). Similarly, the corresponding amino acids in each oligopeptide comprises an L-azidohomoalanine linker. The modified oligonucleotide and the peptide were connected using an azide-alkyne cycloaddition reaction. Four peptide-oligonucleotide conjugates (POC) were formed; each representing one group of key amino acids of the artificial catalytic site (Fig. 2.E-H, S4-S5, and Table S1-S2.). The POCs were modelled together with complementary single stranded (ss) DNA that was later used for the integration of the artificial catalytic site into the DNA nanostructure (see Table S4). The newly formed interactions between the oligopeptides and the oligonucleotides were closely monitored, and *in silico* adjusted to minimize the effect of negatively charged DNA on the emulated catalytic site (Fig. 2.I-J, Fig. S5-S6).

## D-SEPEC assembly

To maintain the relative position and the orientation of the POC elements, a sturdy DNA origami-based hexagonal prism, termed „Shell”, was designed. The Shell comprises two asymmetrical components that were connected by shape-complementary and bridging staple strands (Fig. 3.A, and Fig. S7) [32-34]. The emulated catalytic site was placed in the centre of the Shell structure. The projective lines emanating from the POC oligonucleotides were calculated and used to align the emulated catalytic site towards the optimal position with respect to the inward-facing staple strands of the nanostructure. Single strands complementary to the POC, termed “Carrier Strands”, were placed on these projective lines and merged with the designated staple strand of the Shell (Fig. 3.B, Fig. S7). In this way, the POCs are kept in a stable position relative to each other (via the Carrier Strands) in the central area of the nanostructure with a minimal interaction between the POC elements and the bulk of the Shell and Carrier structures (Fig. 3.C). Once the complete model of the D-SEPEC nanozyme was created, the sequences of the POCs and oligonucleotides were computed and their sequence binding properties were investigated (see Table S4-S5) [35].

**Fig. 3.**
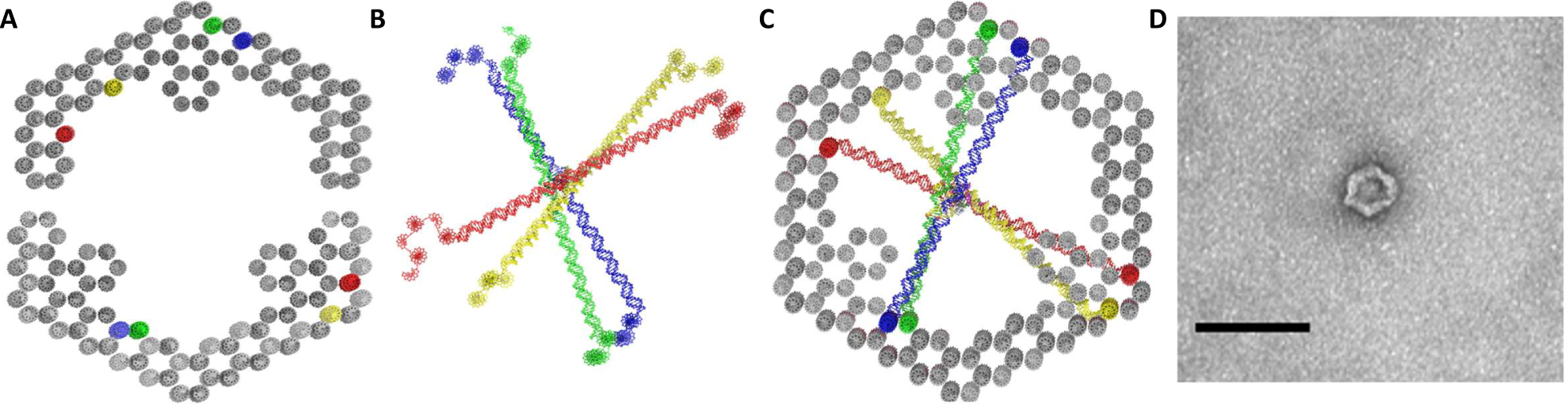
The assembly and integration of the main structural elements of the DNA Scaffold-Embedded Protein Emulation Complex (D-SEPEC). Main structural elements of the D-SEPEC nanozyme and their super assembly. A) Bottom-up representation of the two asymmetrical elements of the hexagonal Shell structure designed to form a stable scaffold for the nanozyme. In the further design process, the interior of the Shell was used as a reference area. The positions in red, yellow, green and blue mark the integration sites of Carrier Strand holding POCs 1 to 4, respectively; B) Following the optimisation of the emulated catalytic site, the short Carrier Strand fragments were linearly extended. The orientation of the elongated Carrier Strands was adjusted so that both ends of each Carrier strand would face the appropriate, inward-facing staple strand of the Shell. Merging of neighboring staple strands and Carrier Strand elements allowed for in silico assembly of the D-SEPEC nanozyme. Colors of Carrier Strands are determined by the associated POC molecules. The position of the emulated catalytic site is marked by the black asterix; C) Top-down representation of the fully assembled D-SEPEC nanozyme. The stability of the DNA-origami Shell structure was employed to maintain the relative orientation of Carrier Strands, and through them the positioning and orientation of the POC elements of the emulated catalytic site. The POC elements are positioned in the intersection area of the Carrier Strands; D) Top-down image of the fully assembled Shell structure as obtained by the negative-staining TEM imaging. Scale bar, 100 nm.

## Structural properties

As confirmed by negative stain TEM imaging (Fig. 3, and Fig. S8), the D-SEPEC nanozyme is a large, rigid, and roughly hexagonally shaped DNA-based nanostructure. The D-SEPEC comprises the Shell (15624 bp), Carrier strands (500bp), and POCs (80 bp linked to 46 amino acids). The molecular weight of the D-SEPEC is approximately 5339 kDa. The height of the hollow Shell structure is ca. 45.8 nm, with the width ranging from 43.4 nm to 46.26 nm. The distance between the emulated catalytic site and the bulk area of the Shell structure was approximately 17.0 nm. The solvent accessible surface area of the nanostructure is approximately 448700 nm^2^, with a peptide area of only 33 nm^2^.

## Catalytic Activity Assay

We performed a series of tests to determine the potential of the D-SEPEC nanozyme to catalyse the exergonic hydrolysis of ATP to ADP and inorganic phosphate (PO_4_^3-^) (Fig. 4.). The phosphate accumulation speed was measured using the Malachite Green Phosphate assay kit [24]. The standardised orthophosphate gradients were used to quantify the turnover rate of the catalytic reactions. All test and control experiments were repeated 110 times, with five incubation intervals of 0, 6, 12, 18, and 24 hours. The final concentration of ATP in each sample type was 0, 25, or 50 µM. In each experiment, test and the control reactions with added ATP were performed in four replicates, while reactions without added ATP were performed in triplicates.

**Fig. 4.**
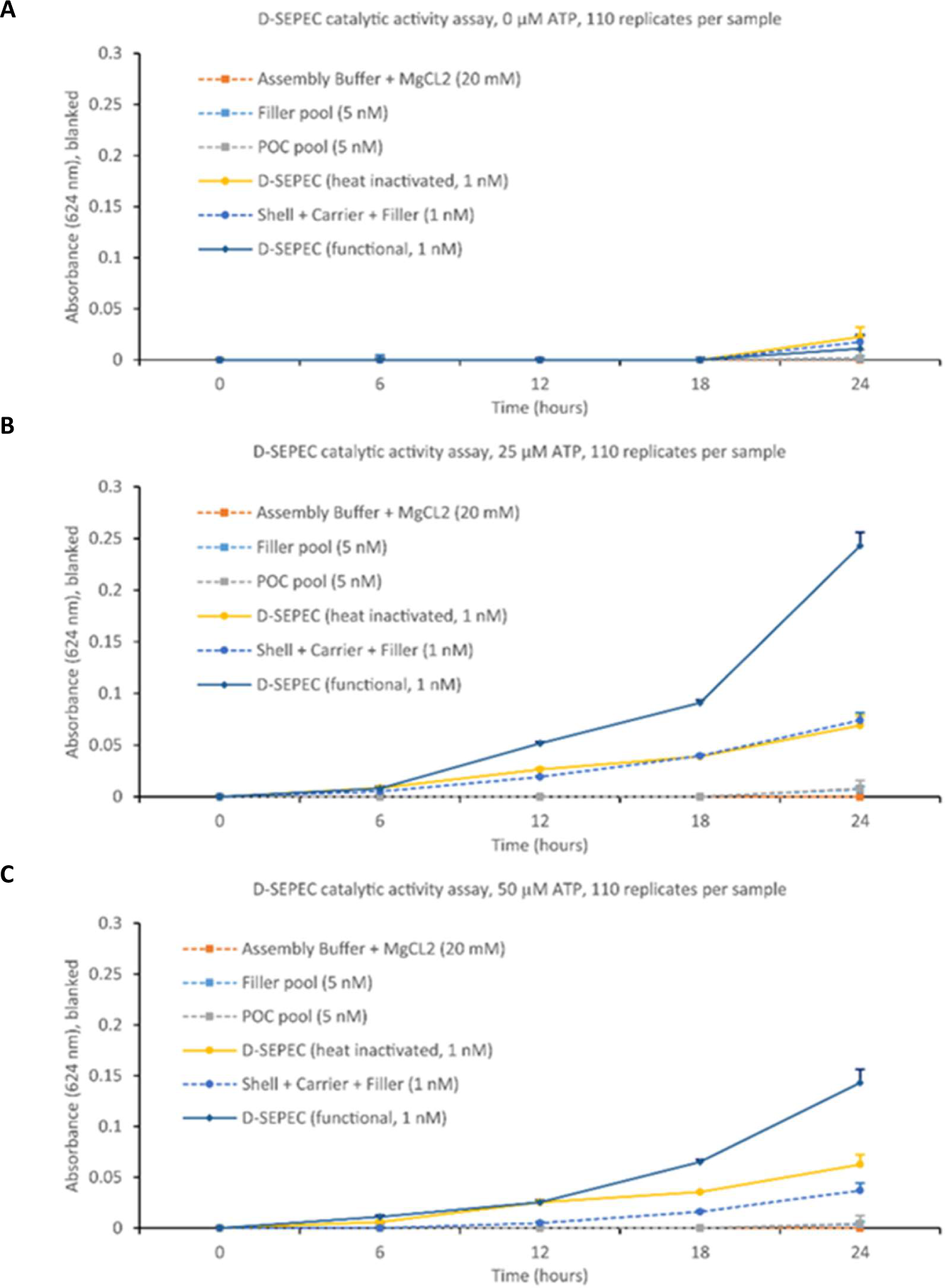
The product generation and substrate turnover measurements. The catalytic activity was measured in discrete time intervals in reactions with 0 (A), 25 µM (B) and 50 µM (C) ATP using the Malachite Green assay. The auto hydrolysis of ATP in the assembly buffer was used to blank the values. The samples with the D-SEPEC nanozyme exhibited the strongest catalytic activity (kcat values of 0.2271 s-1 in systems incubated with 25 µM of ATP, and 0.3622 s-1 in systems with 50 µM of ATP). Heat inactivation of the functional D-SEPEC nanodevice (yellow) is marked by a significant activity drop, close to the levels observed with the non-functional DNA nanostructure (Shell, Carrier and Filler; blue, dotted).

The direct addition of the malachite green reagents to the reaction mixtures resulted in the formation of a visible aggregate, which significantly affected the reliability of the measurements. To solve the aggregation problem, prior to the addition of the malachite reagents a 5 kDa ultrafiltration system was used to purify the resulting phosphate and to regenerate the nanozymes. While the DNA nanostructures and other DNA elements (POCs and controls) remained in a small retention volume, the free orthophosphates, ADP, and ATP were harvested within the flow-through and analysed by high-resolution spectroscopy. Each sample was measured 10 times within several seconds. The data were analysed by comparing the median values of the combined measurements of all replicates of each sample type at the same incubation interval (Fig. 4.A-C).

## Inactivation of the catalytic activity

To confirm that the correct positioning of the POC elements in the D-SEPEC is essential for the catalytic activity, two control experiments were designed. In the first control, only the POC oligonucleotides lacking the peptides (Fillers) were used to construct a non-functional D-SEPEC nanozyme (see Tab. S3). The Fillers were analogously added during the super-assembly process, with the same relative angles as the POCs. The goal of this experiment was two-fold: 1) We wanted to determine if the catalytic properties of the nanozyme originated from the peptides in the D-SEPEC, or from randomly occurring DNAzymes in the staple strands and/or Shell. 2) By creating a structure entirely made of DNA, we wanted to measure the effect that the natural degradation and/or possible DNAzyme sequences inside of the large DNA-based nanostructure might have on the effectiveness and the resolution of the malachite green assay (see Fig. 4, Shell-Carriers-Fillers).

In the second control, the functional D-SEPEC nanozymes were incubated at 95 °C for 15 minutes. This resulted in the denaturation of their structure and the disassembly of the D-SEPECs. After cooling to room temperature, the evaluation of the catalytic activity was initiated by addition of the substrate and incubation for up to 24 hours. The goal of this experiment was to verify that the catalytic activity originates from the correct and predefined assembly, and not from random and unspecific interaction between the DNA and POC molecules (Fig. 4).

## Substrate turnover analysis

Based on the substrate turnover in different sample types and controls, several observations have been made. In the control samples with no ATP, during the incubation period of 24 hours, the absorbance of the malachite green reagent was minimally changed. Only a slight increase was observed in samples containing the Shell structure, which can be attributed to the slow degradation of the DNA-based nanostructures (Fig 4.A). In reactions with 25 µM ATP, a slow increase of the PO_4_^3-^ concentration in buffer control samples indicated a natural degradation of ATP that occurs under the given conditions (Fig 4.B) [28]. As such, the buffer control measurements were used as a baseline in determining the substrate turnover speed. A low turnover rate was also observed in the control samples (POC and Filler pools), showing that the random combinations of POC molecules are unable to catalyse the hydrolysis of ATP. Furthermore, this observation confirmed that the Filler molecules have no DNAzyme activity (Fig 4.C).

In control reactions with the correctly assembled but non-functional nanozymes (made from Shell, Carrier and Filler elements), we observed significantly lower turnover rates (k_cat_ = 0.0567– 0.1361 s^-1^) compared to the samples containing the functional D-SEPEC nanozymes (k_cat_ = 0.2271 – 0.3622 s^-1^) (Fig. 4.A-C). The product accumulation in the samples containing the non-functional nanozymes indicated that our DNA-based superstructure exhibits a low catalytic effect on the hydrolysis of ATP substrate. Due to the high amount of the staple strands in the nanostructure, we could not exclude the possibility that some of them might possess the capacity to hydrolyse ATP. The heat inactivated D-SEPEC and the non-functional nanozyme exhibited similar turnover rates (Fig. 3.B).

Heat denaturation was successfully used to destabilise the correctly assembled, functional D-SEPEC nanozyme. By heating the samples to 95 °C and cooling them to room temperature prior to the addition of the substrate, we significantly hindered the generation of the product in the reactions (k_cat_ = 0.1114– 0.1253 s^-1^). This suggests that the POC elements of the active centre can hydrolase ATP only if correctly assembled. The unspecific aggregation of the structures led to product generation levels similar to those observed in the Shell-Carrier-Filler reactions.

The catalytic properties of D-SEPEC nanozymes built in three separate (from start to finish) assembly processes were analysed in a total of 1330 reactions, spread over sample types named D-SEPEC 1, 2, and 3 (Fig 4.A-C, and Fig. S9). In comparison to the controls, the D-SEPEC nanozymes consistently exhibited the highest catalytic activity. We calculated the turnover rate of the D-SEPEC artificial catalytic site (k_cat_ = 0.2271 – 0.3622 s^-1^) to be in the range of 10 to 20 ATP molecules per D-SEPEC and per minute of incubation at RT (Fig. 4). Recent studies on the isolated *Sulfolobus’s* FlaI reported the turnover rate of the enzyme to be in the range of the D-SEPEC nanozyme, with approximately one ATP per enzyme per minute [3]. A similar turnover rate was observed in the reactions where 50 µM ATP were used for the incubation (Fig. 4). The increase of the substrate concentration led to a higher coherence of measurements between the different D-SEPEC sample pools that were separately assembled.

## Discussion

Industrial and biomedical exploitation of enzymatic catalysis is limited by the currently rather poor ability to emulate catalytic properties in artificial environments. In this study, we demonstrate a *de novo*, bottom-up approach for the development of a DNA-based nanozyme able to catalyse ATP hydrolysis. Previous artificial enzyme strategies were predominantly based on a small number of amino acids or functional groups usually integrated directly into an inert and stable protein scaffold structure or matrix [7, 12-13, 18-19]. Rothlisberger et al. (2008), presented an approach for a bottom-up design of a protein capable of catalysing the Kemp elimination reaction. In their work, key amino acids of an idealised active site were encoded into fitting structural domains of proteins, primarily TIM barrels. Although successful in achieving substantial catalytic rates (k_cat_=0.018 – 1.37 s^-1^), the authors suggested that the rigidity of the construct, as a main shortcoming of their design, weakens the substrate and transition-state binding affinity. A similar issue, with significantly lower catalytic rates (k_cat_≈2 x 10^-4^), was reported in studies on molecular imprinting of the key functional groups in a polymer matrix [12-13]. It is noteworthy that the positioning of the functional groups in the matrix was determined exactly by their interactions with the transition state analogue. The authors argue that the number of key residues that can be embedded determines the catalytic performance of the artificial enzymes. The positioning of peptides is challenging, and these previously published strategies are not suitable for the *de novo* design of catalytic sites with secondary or tertiary motifs. For this reason, we designed a system that can expand the emulated catalytic site to include and organise residues indirectly involved in the idealised catalytic mechanism, and to shape them into the structural motifs of enzymes.

In our design approach, the key residue’s count is only limited by the ability to synthesise POC molecules of a certain length. Through interactions with a relatively stable framework structure, modular elements organise secondary structures of POCs into a tertiary structure of the emulated catalytic site. Significance of this can be found in the work of Romero et al. (2018), where sequence and structural alignments of proteins were used for the creation of a structural motif’s catalogue involved in the active site of NTPases, foremost Walker-A sequences. The evolutionary conserved motifs were separately used in a *de novo* design of self-folding peptides [18]. The observed low catalytic activity (k_cat_= 5.5 x 10^-4^ s^-1^) was attributed to the inability of the approach to assemble several motifs into a tertiary structure, which resulted in a lack of the magnesium-binding site and the active-site cavity.

In contrast to the Shell and Carrier Strands, the catalytic centre of the D-SEPEC was designed to be a conformationally flexible structure, stabilised by the non-covalent interactions of the POC molecules and their environment. The design principle was based on the premise that the structure as a whole will be close to the minimum of its energy potential if all elements of the structure are close to their steric and electrostatic minimums. As the introduction of stable structural elements can lead to the stabilisation of the whole structure, the quality of the structural, phylogenetic, and kinetic data, together with the extent of the molecular dynamics simulations of the native protein, were important aspects during the design process. During the simulations, the D-SEPEC resembled the catalytic structure only in a certain percentage of time, inversely proportional to the energy of the modelled structure. Nevertheless, the precise *in silico* positioning of the POC molecules is essential, as demonstrated in the control experiments (Fig. 3).

The substrate turnover rates of the D-SEPEC nanozyme and the native enzyme exhibited similar levels. The *Sulfolobus’s* FlaI is a relatively slow ATPase with a turnover rate of up to two orders of magnitude lower than some of its homologues [23]. For our calculations of the turnover rate, we assumed the concentration of the nanostructures in each reaction being the maximally possible concentration for measured amount of DNA. However, based on the general experience in the DNA nanotechnology field, it is reasonable to assume that the real concentration of the correctly assembled nanozymes is significantly lower. Thus, the lower percentage of the properly assembled nanostructures means an equally higher turnover rate of the individual unit.

In conclusion, we have demonstrated a novel approach to use the essential elements of the protein’s active site, as a starting point to design a highly precise, DNA-based nanozyme capable of *in situ* and *de novo* recreation of the original catalytic properties. We have successfully recycled the nanozymes in multiple experiments. In contrast to proteins, an outstanding benefit of this technology is that the DNA-based self-assembly process allows a simple thermal regeneration process if the artificial enzymes get, e.g., heat denatured. Chaperons to refold the nanozyme are not required. We observed that the correct assembly of the D-SEPEC plays a major role in the emulation of the catalytic activity, as only correctly positioned POC molecules would maintain the turnover speed of the reaction. By taking into account the turnover speed (k_cat_ = 0.2271 – 0.3622 s^-1^) observed in the reactions of the functional D-SEPEC nanostructures, which is *on par* with the native FlaI enzyme, we can conclude that further improvements to this technology would lead to substantial advances in the fields of artificial enzymes and nanotechnology. Future designs will be significantly facilitated by a novel molecular modelling software tool that was developed in parallel by our group [38]. Using this approach for enzymes of extremely conserved secondary structures and rapid turnover rates could be beneficial for the development of, e.g., new diagnostic tools, while the functional emulation of slow enzymatic reactions or hard to purify enzymes could open new possibilities in the fields of chemical industry.

## Materials and methods

The design and experimental procedures are detailed in the Supplementary Materials.

### Self-assembly and purification of the D-SEPEC nanozyme

The fabrication of D-SEPEC nanozymes was carried out in sequence, following the accepted guidelines on the assembly of DNA-origami based nanostructures [27,28]. The self-assembly was conducted in an assembly buffer, under neutral pH and high salt environment (50 mM TRIS buffer, 18 mM MgCl_2_, pH=7.0). The folding reactions of each Shell element were PEG purified by the method reported by Stahl et al. (2014) using a PEG precipitation buffer (15% (w/v) PEG-8000, 5 mM Tris, 505 mM NaCl) [28]. All oligonucleotide sequences were synthesized by IDT BVBA (Leuven, Belgium). POC molecules were synthesized by Biomers.net GmbH (Ulm, Germany). To avoid any ATPase activity by enzymatic contaminants, all reagents were heat inactivated at 95° for 15 min.

### Negative staining transmission electron microscopy (nsTEM)

The morphological examination of nanostructures was done using transmission electron microscope (TEM) performed on a Morgagni operated at 80 kV. The images were taken by a Morada camera. Samples were prepared by dropping 5 µl of diluted nanostructures on glow-discharged carbon-coated Cu400 TEM grids and negative staining using 2% uranyl acetate (see Fig. S8).

### Catalytic activity assay and product separation

Test and control reaction samples were subjected to a range of substrate concentrations (0, 25, and 50 uM ATP) and incubation times (0, 6, 12, 18, and 24h). The concentration of all nanostructures involved in the experiments was 1 nM. The concentration of the POC and Filler pool elements was 5 nM. Following the incubation, the generated products were separated from the nanostructures on ultrafiltration columns (15 000 g, 15 min, 5kDa pores, Amicon). The flow-through samples were transferred to black, non-binding, clear bottom, plate wells containing malachite green reagent (Greiner Bio-one, fisher scientific). Samples were quickly mixed and measured by high resolution spectroscopy (EnSpire, PerkinElmer). The measured wavelength was 624 nm, with each sample measured 10 times over several seconds.

### Recycling of the nanostructures

The DNA nanostructures retained in the ultrafiltration reservoir were immediately reused. By ultra-filtrating each sample in 500 µ1 of the assembly buffer, the leftover traces of ATP, ADP and PO_4_^3-^ were removed. To initiate the next experiment, samples were diluted to their original concentration, using the 50 µl of the assembly buffer (50 mM TRIS buffer, 18 mM MgCl2, pH=7.0), and the appropriate amounts of the substrate were added.

## ACKNOWLEDGMENTS

We thank J. Angerer and R. Soldo for technical assistance, and S. V. Albers, J. L. Anderson, B. Bertosa, D. Rakoczy and T. Miletic for discussions. This project has received funding from the European Union’s Horizon 2020 research and innovation program under grant agreement No. 686647. T. Kekic wrote the manuscript, developed the D-SEPEC in silico and performed the functionality experiments. Y. Ahmadi reviewed the manuscript, developed the DNA nanostructure and made the nsTEM images. I. Barisic designed the study and wrote the manuscript. Authors declare no competing interests. All data is available in the main text or the supplementary materials. Raw data from in silico analysis and colorimetric measurements, as well as detailed schematics and sequences of the DNA-origami elements are available from the corresponding author on request.

## SUPPLEMENTARY MATERIALS

Materials and Methods

Figs. S1 to S10

Tables S1 to S5

References (39-46)

## REFERENCES AND NOTES

1. F. Berger et al., Regulation of poly(ADP-ribose) polymerase 1 activity by the phosphorylation state of the nuclear NAD biosynthetic enzyme NMN adenylyl transferase 1. Proc. Natl. Acad. Sci 104, 3765–70 (2007). doi: 10.1073/pnas.0609211104

2. X. Chang et al., Role of N-linked glycosylation in the enzymatic properties of a thermophilic GH 10 xylanase from Aspergillus fumigatus expressed in Pichia pastoris. PLOS One (2017). doi: 10.1371/journal.pone.0171111

3. S. Reindl et al., Insights into FlaI Functions in Archaeal Motor Assembly and Motility from Structures, Conformations, and Genetics. Mol. Cell. 49, 1069–82 (2013). doi: 10.1016/j.molcel.2013.01.014

4. Amit et al., An antioxidant nanozyme that uncovers the cytoprotective potential of vanadia nanowires. Nature Commun. 5, 5301 (2014). doi: 10.1038/ncomms6301

5. Linko, V. et al., A modular DNA origami-based enzyme cascade nanoreactor. Chem. Commun. 51, 5351–5354 (2015). doi: 10.1039/C4CC08472A

6. M. Liu et al., A DNA tweezer-actuated enzyme nanoreactor. Nature Commun. 4, 2127 (2013). doi: 10.1038/ncomms3127

7. D. Rothlisberger et al., Kemp elimination catalysts by computational enzyme design. Nature 453, 190–195 (2008). doi: 10.1038/nature06879

8. D.W. Watkins et al., Construction and in vivo assembly of a catalytically proficient and hyperthermostable de novo enzyme. Nature Commun. 8, 358 (2017). doi: 10.1038/s41467-017-00541-4

9. E. Donnelly et al., A de novo enzyme catalyzes a life-sustaining reaction in Escherichia coli. Nat. Chem. Biol. 14, 253–255 (2018). doi: 10.1038/nchembio.2550

10. G. Grossi et al., Control of enzyme reactions by a reconfigurable DNA nanovault. Nature Commun. 8, 992 (2017). doi: 10.1038/s41467-017-01072-8

11. A. Katz and M. E. Davis, Molecular imprinting of bulk, microporous silica. Nature 403, 286–289 (2000). doi: 10.1038/35002032

12. J. Liu and G. Wulff, Molecularly Imprinted Polymers with Strong Carboxypeptidase A-Like Activity: Combination of an Amidinium Function with a Zinc-Ion Binding Site in Transition-State Imprinted Cavities. Angew. Chem. Int. Ed. 43, 1287–90 (2004). doi: 10.1002/anie.200352770

13. C.V. Hanson et al., Catalytic antibodies and their applications. Curr. Opin. Biotech. 16, 631–36 (2005). doi: 10.1016/j.copbio.2005.10.003

14. F. Richter et al., De Novo Enzyme Design Using Rosetta3. PLOS One (2011). doi: 10.1371/journal.pone.0019230

15. O. I. Wilner et al., Enzyme cascades activated on topologically programmed DNA scaffolds. Nature Nanotech. 4, 249–254 (2009). doi: doi.org/10.1038/nnano.2009.50

16. S. Walsh et al., DNA Cage Delivery to Mammalian Cells. ACS Nano 5, 5427–32 (2011). doi: 10.1021/nn2005574

17. Y. Zhang et al., Proximity does not contribute to activity enhancement in the glucose oxidase– horseradish peroxidase cascade. Nature Commun. 7, 13982 (2016), doi: 10.1038/ncomms13982

18. M. L. Romero et al., Simple yet functional phosphate-loop proteins. PNAS 115, 11943–50 (2018). doi: 10.1073/pnas.1812400115

19. M. D. Arifuzzaman, Y. Zhao, Artificial Zinc Enzymes with Fine-Tuned Catalytic Active Sites for Highly Selective Hydrolysis of Activated Esters. ACS Catal. 8, 8154–8161 (2018). doi: 10.1021/acscatal.8b02292

20. Y. Choi et al., DNA Origami-Based Förster Resonance Energy-Transfer Nanoarrays and Their Application as Ratiometric Sensors. ACS Appl.Mater. Interfaces 10, 23295–23302 (2018). doi: 10.1021/acsami.8b03585

21. S. M. Douglas et al., A Logic-Gated Nanorobot for Targeted Transport of Molecular Payloads. Science 335, 831–834 (2012). doi: 10.1126/science.1214081

22. D. Han et al., DNA Origami with Complex Curvatures in Three-Dimensional Space. Science 332, 342–346 (2011). doi: 10.1126/science.1202998

23. P. Chaudhury et al., Characterization of the ATPase FlaI of the motor complex of the Pyrococcus furiosus archaellum and its interactions between the ATP-binding protein FlaH. Peer J. 6, 4984 (2018). doi: 10.7717/peerj.4984

24. B. Repen et al., Optimization of a malachite green assay for detection of ATP hydrolysis by solubilized membrane proteins. Anal. Biochem. 15, 103–5 (2012). doi: 10.1016/j.ab.2012.04.016

25. E. Marras et al., Programmable motion of DNA origami mechanisms. PNAS 112, 713–718 (2015). doi: 10.1073/pnas.1408869112

26. P. Kosuri et al., Rotation tracking of genome-processing enzymes using DNA origami rotors. Nature 572, 136 –140 (2019). doi: 10.1038/s41586-019-1397-7

27. F. A. Kiani, S. Fischer, Comparing the catalytic strategy of ATP hydrolysis in biomolecular motors. Phys. Chem. Chem. Phys. 18, 20219–233 (2016). doi: 10.1039/C6CP01364C

28. F. Praetorius, H. Dietz, Self-assembly of genetically encoded DNA-protein hybrid nanoscale shapes. Science 355, 5488 (2017). doi: 10.1126/science.aam5488

29. C. B. Harrison, K. Schulten, Quantum and classical dynamics simulations of ATP hydrolysis in solution. J. Chem. Theory. Comput. 8, 2328–2335 (2012). doi: 10.1021/ct200886j

30. E. Stahl et al., Facile and scalable preparation of pure and dense DNA origami solutions. Angew. Chem. Int. Ed. 53, 12735–40 (2014). doi: 10.1002/anie.201405991

31. M. Priess et al., Molecular Mechanism of ATP Hydrolysis in an ABC Transporter. ACS Cent. Sci. 4, 1334–1343 (2018). doi: 10.1021/acscentsci.8b00369

32. M. Dittrich et al., On the Mechanism of ATP Hydrolysis in F1-ATPase. Biophys. J. 85, 2253–2266 (2003). doi: 10.1016/S0006-3495(03)74650-5

33. Koga N, et al. Principles for designing ideal protein structures. Nature 491, 222–227 (2012). doi: 10.1038/nature11600

34. S. M. Douglas et al., Rapid prototyping of 3D DNA-origami shapes with caDNAno. Nucleic Acids Res. 37, 5001–6 (2009). doi: 10.1093/nar/gkp436

35. D. N. Kim et al., Quantitative prediction of 3D solution shape and flexibility of nucleic acid nanostructures. Nucleic Acids Res. 40, 2862–8 (2012). doi: 10.1093/nar/gkr1173

36. R. Veneziano et al., Designer nanoscale DNA assemblies programmed from the top down. Science 352, 1534 (2016). doi: 10.1126/science.aaf4388

37. R. Owczarzy et al., IDT SciTools:a suite for analysis and design of nucleic acid oligomers. Nucleic Acids Res. 36, 163–169 (2008). doi: 10.1093/nar/gkn198

38. E. Liano et al., Adenita: Interactive 3D modeling and visualization of DNA Nanostructures. BioRxiv (2019). doi: 10.1101/849976

