## Supplementary materials for "An Enzymatic Active Site Embedded in a DNA Nanostructure"

**This PDF file includes:**

Materials and Methods
Figs. S1 to S10
Tables S1 to S5
References (*39-46*)

### Materials and Methods

Inoculation, expression and purification of p7560 and p8064 ssDNA scaffolds

Cultures of the XL-1 blue E.coli were formed by addition of 1.5 ml of overnight (ON) bacterial cells to the pre-warm flask (CorningTM Erlenmeyer baffled cell culture flasks, 500 ml) including 125 ml of Low salt LB media. Cultures were incubated at 37 °C until OD600 was in range of 0.3 – 0.6. Separate cultures were inoculated by addition of 50 ul of p7560 or p8064 bacteriophages and shake-incubated at 37 °C at 200 rpm for 5 hours. To pellet the bacteria, cultures were transferred to the fresh containers (CorningTM Falcon Conical Centrifuge Tubes, 50 ml) and centrifuged at 4000 g at 4 °C. The supernatant was transferred to the fresh containers (CorningTM Falcon Conical Centrifuge Tubes, 50 ml) and 9 ml of Phage Precipitation buffer was added (2.5 M NaCl, 20% PEG8000 (w/v). Samples of phages were incubated in ice-water bath overnight. Following the incubation, to pellet the phages, samples were centrifuged at 16 000 g for 15 minutes at 4 °C. The supernatant was discarded and samples were solubilised in 1.9 ml of Phage Resuspension buffer (10 mM Tris, 1 mM EDTA, pH=8.5).

Two volumes of Lysis buffer (0.2 M NaOH, 1% SDS) was added to resuspended phages and gently mixed by inversion on room temperature (RT). Further, 1.5 volume of Neutralisation buffer (3 M KOAc, pH=5.5) was added to the samples and gently mixed by inversion on RT. Samples were incubated in ice-water bath for 15 minutes followed by the centrifugation at 16 000 g for 15 minutes at 4 °C. Supernatant was carefully discarded. Pellets were gently triturated in 1.5 ml of 75 % ethanol and transferred to 2 ml tubes (EppendorfTM Safe-Lock Tubes). Samples were incubated in ice-water bath for 10 minutes. Following the incubation, samples were centrifuged at 16 000 g for 15 minutes at 4 °C. Supernatant was carefully discarded and the pellets were dried for 10 minutes. Dried pellets were resuspended in 200 μl of the Scaffold Resuspension buffer (50 mM Tris, pH=7.0), followed by the Agarose gel electrophoresis (AGE) and spectroscopic measurement of DNA concentration.

Self-assembly and PEG purification of the Sp7560 and Sp8064 elements of the Shell structure

Two structural elements of the Shell structure named Sp7560 and Sp8064 (Shell-plasmid7560, Shell-plasmid8064) were self-assembled in the separate reactions following the accepted guidelines on the assembly of DNA-origami nanostructures [28,30]. The reaction mixture was created by addition of scaffold plasmid and associated staple strands in the 1 : 10 ratio to the Assembly buffer (50 mM Tris, 5 mM NaCl, pH=7.0) containing 18 mM MgCl2. In each assembly reaction the final concentration was 20 and 200 nM for scaffold and staple strands, respectively. The extended staple strands used for integration of the Carrier Strands were added in higher concentration (400 nM). Annealing was done by exposing the reactions to 95°C for 15 minutes, followed by the prolonged cooling incubation from 94 °C to 25 °C with temperature gradient of 1 °C every 90 minutes.

The nanostructures were purified using the purification method reported by Stahl et al.[30]. In brief, 100 ul of DNA origami samples in Assembly buffer (50 mM Tris, 5 mM NaCl, pH=7.0) including 18 mM MgCl2 were mixed with the equivalent volume of 22 mM MgCl2 supplemented Assembly buffer followed by the addition of 200 ul of PEG precipitation buffer (15% (w/v) PEG 8000, 50 mM Tris and 505 mM NaCl). The solution was then mixed gently by tube inversion and spinned at 16 000 g at R.T. for 25 minutes. The supernatant was carefully discarded and the pellet dissolved in 20 ul of Assembly buffer and incubated for 1 day at R.T. with mild agitation (60 rpm). Incubation was followed by the spectroscopic measurement of DNA concentration and AGE.

Super-assembly of the D-SEPEC nanozyme

The super-assembly reactions of the D-SEPEC nanozyme and inactive Shell-Carriers-Fillers control nanostructure was done by pooling the equimolar concentrations of preassembled Shell elements with five times higher concentration of Carrier Strands and POC/Filler elements, in Assembly buffer (50 mM Tris, 5 mM NaCl, pH=7.0) with 18 mM MgCl2. The final concentration of Sp7560 and Sp8064 elements were 15 nM per reaction. Super-assembly was done by incubating the reactions on 60°C for 15 minutes, followed by the prolonged cooling incubation from 59 °C to 25 °C with temperature gradient of 1 °C every 90 minutes. Following the assembly spectroscopic measurement of DNA concentration and AGE were conducted.

Molarity measurement of the nanostructures

Samples of 2 ul of each reaction mixtures were taken in three replicates, and spectroscopically measured by using the NanoDrop 2000 (NanoDropTM, Theromo Scientific). NanoDrop 2000 software package was used to calculate the mass concentration and purity of DNA in samples. Maximum possible molarity of the nanostructures was calculated by dividing the mass concentration of samples with molar mass of nanostructure in the reaction mixtures. Molar mass was calculated by multiplying the number of base pairs of the modelled nanostructure with the average molar mass of a single base pair (330 g / mol).

Negative staining transmission electron microscopy (nsTEM)

The morphological examination of nanostructures was done using transmission electron microscope (TEM) performed on a Morgagni operated at 80 kV. The images were taken by a Morada camera. Samples were prepared by dropping 5 µl of diluted nanostructures on glow-discharged carbon-coated Cu400 TEM grid and negative staining using 2% Uranyl acetate (Fig. S5,b).

Catalytic activity assay and product separation

Test and control reaction samples were subjected to the range of substrate concentration (0, 25 and 50 uM ATP) and incubation times (0, 6, 12, 18 and 24 hours) under RT. The reaction buffers in all samples were maintained under the neutral pH and high salt conditions (50 mM TRIS buffer, 20 mM MgCl2, pH=7.0). Concentration of all nanostructures involved in the experiments was 1 nM. Concentration of POC and Filler pools elements was 5 nM. Following the incubation, generated product was separated from the nanostructures on the ultrafiltration columns (15 000 g, 15 min, 5kDa pores, Amicon). The flow through samples were transferred to the black, non-binding, clear bottom, plate wells (Greiner Bio-one, fisher scientific). One volume of the malachite green reagent was added to four volumes of each samples (Malachite Green Phosphate Assay Kit, Sigma-Aldrich). The samples were quickly mixed and measured by the high resolution spectroscopy (EnSpire, PerkinElmer). Measured wavelength was 624 nm, with each sample measured 10 times over several seconds.

Recycling of the nanostructures

The DNA nanostructures retained in the ultrafiltration reservoir was immediately reused. By ultra-filtrating each sample in 500 µl of the Assembly buffer (15 000 g, 15 min, 5kDa pores, Amicon), the leftover traces of ATP, ADP and PO43 were removed. To initiate the next experiment samples were diluted to their original concentration, using the 50 ul of the Assembly buffer (50 mM TRIS buffer, 18 mM MgCl2, pH=7.0) with added correct amounts of the substrate.

Molecular dynamics simulations of the FlaI protein complex

Domain dynamics and the interactions between the enzyme's active site, the substrate, the products and the cofactor were analyzed by using the molecular dynamics (MD) simulations. The crystal structure of the FlaI hexamer (PDB: 4IHQ) was used to generate the protein complex systems [40]. The systems of approximately 290 000 atoms were built and simulated using GROMACS 5.1.4 and AMBER 16 program packages [3,39]. Water molecules found in the crystal structure within 0.3 nm from the protein heavy atoms were retained for calculation. The hydrogen atoms were added using the Tleap module of the AMBER 16 program package. The proteins were placed in the center of the rectangular parallelepiped system with implementation of the periodic boundary conditions (PBC). The TIP3P water molecules were added as a solvent. In order to neutralize the system, randomly chosen solvent molecules were replaced with the Na+/Cl- ions. Substrates used in the separate MD simulations were ATP, ADP with PO4 and ADP. For parametrization of the protein atoms, the AMBER ff99SB force field was used. Parameters for ADP, ATP, PO4 and Mg ions were obtained using the AMBER parameter database [39 – 41]. All systems were energy minimized and their geometry optimized in a process consisting of 50000 steps of the steepest descent algorithm. The constraint of 418.4 Kj was applied on the protein complex atoms, while all the solvent molecules remained unconstrained. The temperature was linearly increased from 0 to 300 K during the first 250 ps, followed by 50 ps of simulation at a constant temperature (300 K) using the Berendsen thermostat. During the first 600 ps, protein complex atoms were constrained with the force constant of 104.6 Kj and the volume was kept constant. From 600 ps to the end of simulations, no constraints were applied and the simulations were conducted at a constant temperature (300K) and pressure (101 325 Pa) using the Berendsen thermostat and barostat. The time step was 1 fs, and the structures were sampled every 5 picosecond. The periodic boundary conditions (PBC) were applied. Particle mesh Ewald (PME) was used for the calculation of the electrostatic interactions. The cut-off value for the non-bonded interactions was set to 1 nm. Following the geometry optimization, the systems were subjected to MD simulations with the final trajectory length of 200ns.

Trajectories were explored globally, on the level of protein complex, by measuring the Root-mean-square-deviations (RMSD), gyration radius (Rg), Solvent accessible surface (SAS) and relative domain deviations. On the level of residues, the analysis was based on the non-covalent interactions of the active site of the protein, and the comparison of the binding profiles between substrates/product and active site's observed throughout the trajectories. The structure of the active site of the FlaI, obtained from MD simulations, was further analysed by the structural comparison with the active sites of series of ATPases (PDB: 4II7, 4IHQ, 4RVC, 4YMV, 2YZ2, 1G6H, 169X, 1GAJ, 2BJW, 5Z6R, 1LV7, 2ZAN, 5C18, 2QZ4). All systems were simulated and analyzed using the GROMACS, AMBER, SCHRODINGER and VMD program packages [3,39,42,43].

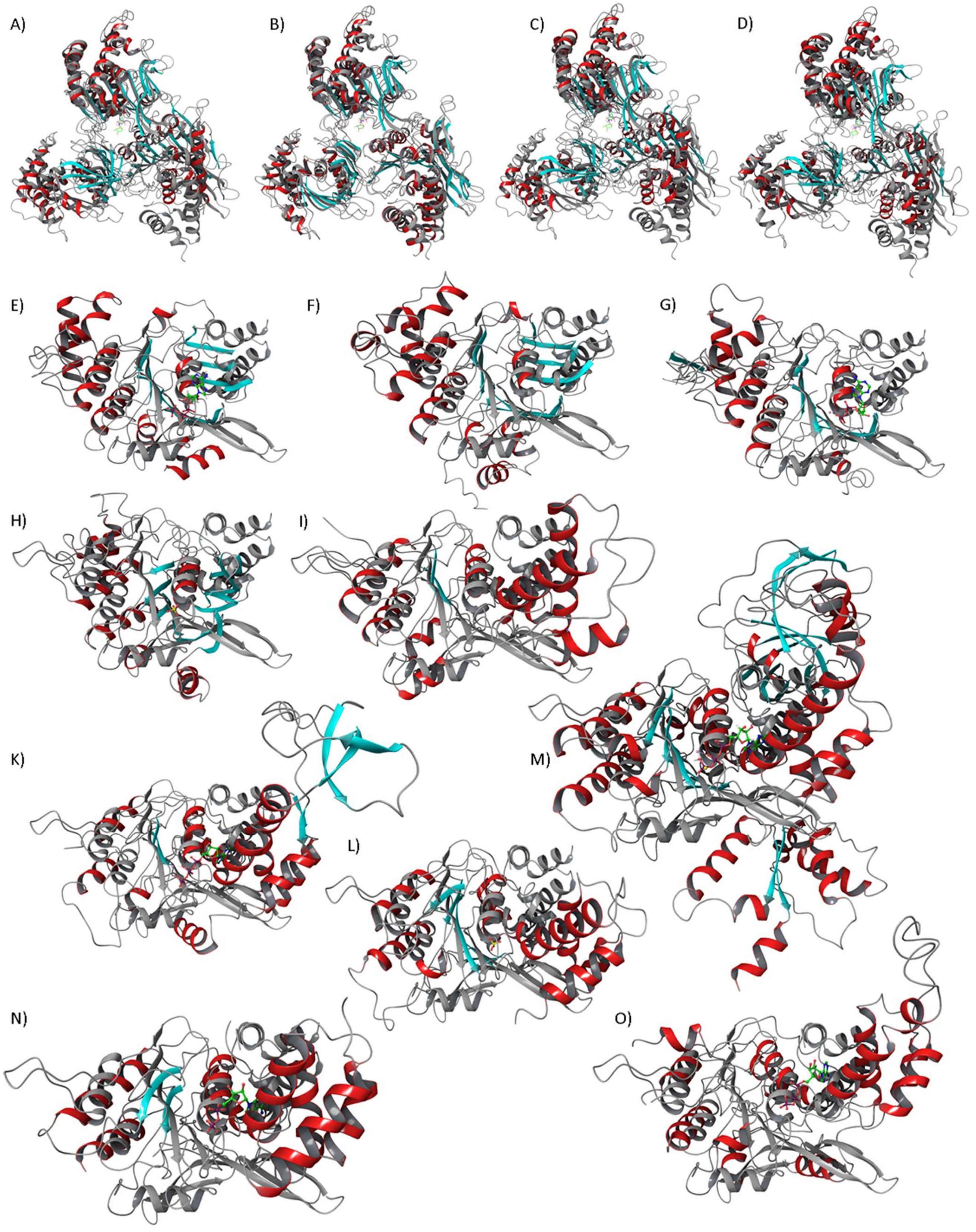
A)
B)
C)
D)
E)
F)
G)
H)
I)
K)
M)
L)
N)
O)

**Fig. S1. The structural homology analysis of the FlaI protein complex.**

The representatives of the p-loop NTPase domain superfamily structurally aligned with the active site of the FlaI ATPase. The structure of the FlaI, based on the crystal structure (PDB:4IHQ), was obtained after 200 ns of MD simulation. Structure obtained from the simulation was compared, first with the conformational states of FlaI monomers found in the crystal structure 4IHQ (A-C), followed by comparison with the structures under PDB code: 4II7 (D), 4RVC (E), 4YMV (F), 2YZ2 (G), 1G6H, 169X (H), 1GAJ (I), 2BJW (J), 5Z6R (K), 1LV7 (L), 5C18 (M), 2ZAN (N), 2QZ4 (O). Alpha helices and beta sheets are coloured in red and blue respectively, while reference structure is coloured in gray.

A)

B)

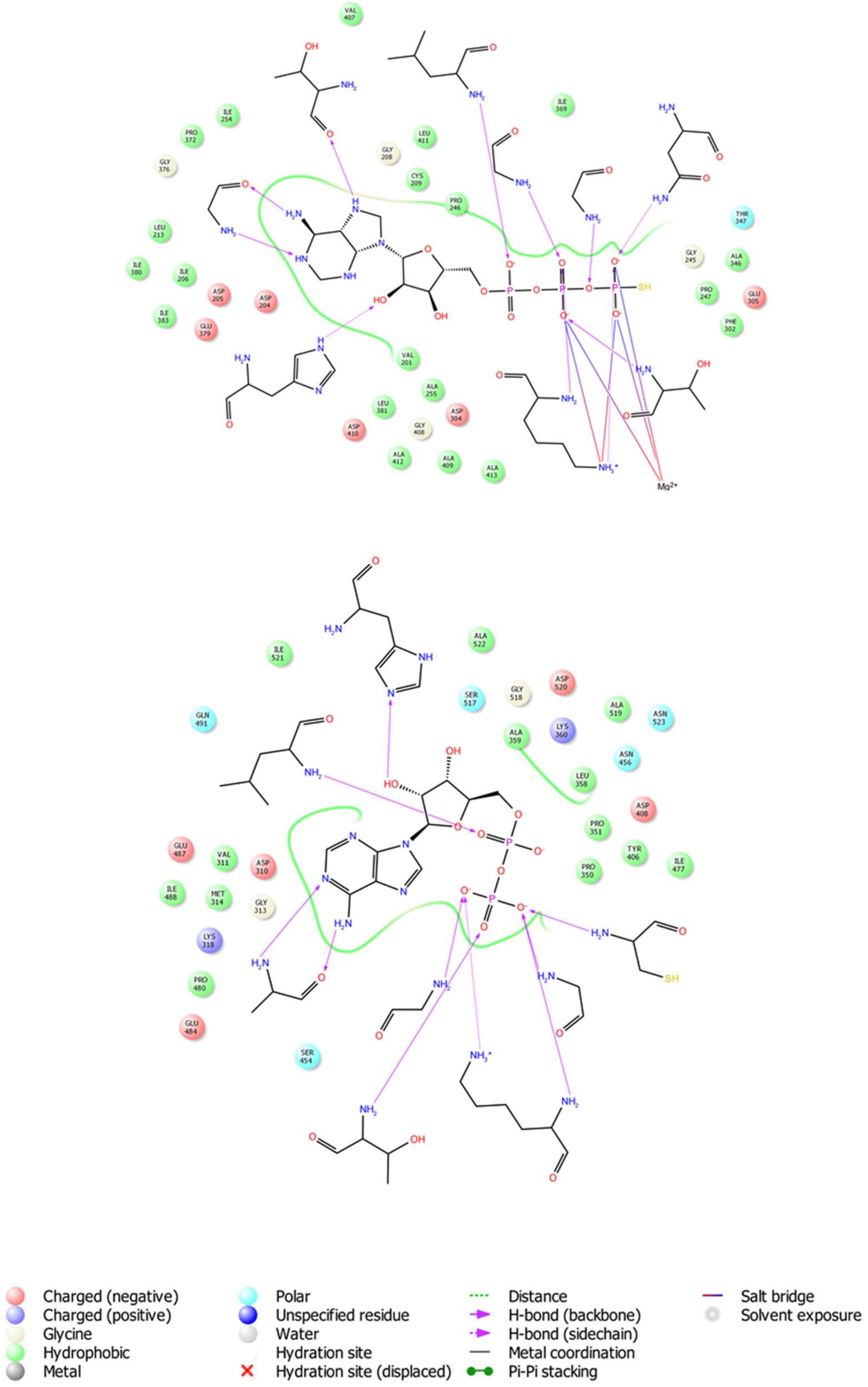

Charged (negative)
Charged (positive)
Glycine
Hydrophobic
Metal
Polar
Unspecified residue
Water
Hydration site
Hydration site (displaced)
Distance
H-bond (backbone)
H-bond (sidechain)
Metal coordination
Pi-Pi stacking
Salt bridge
Solvent exposure

C)

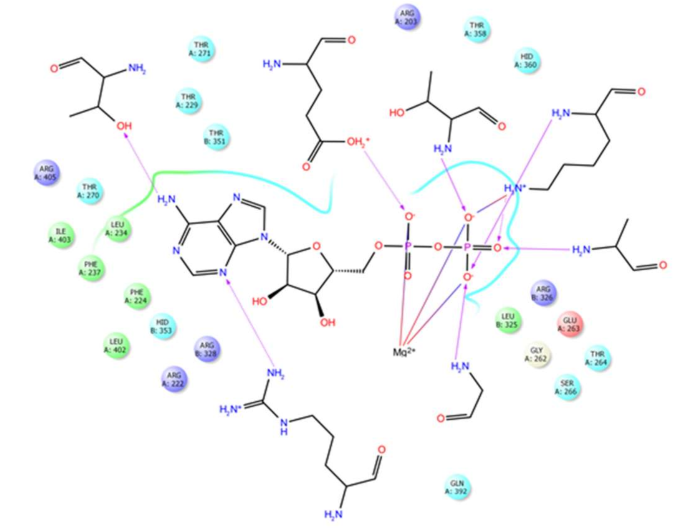

D)

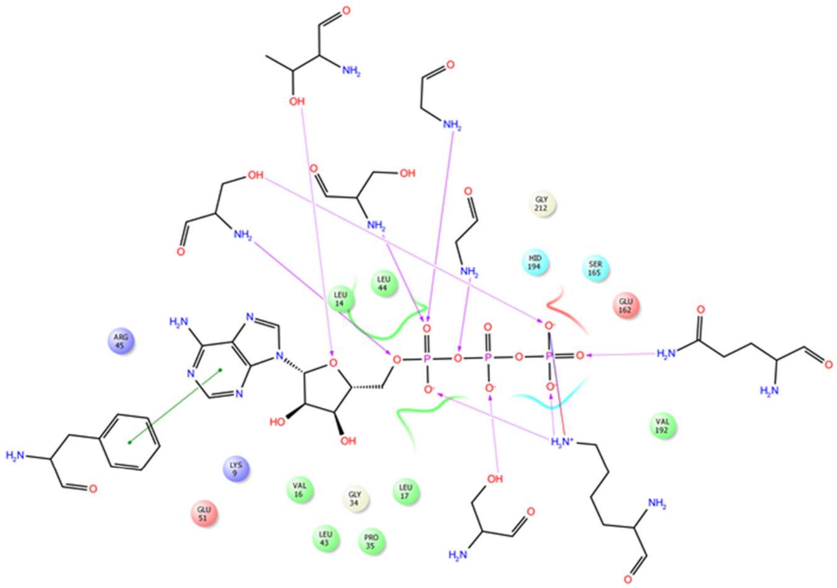

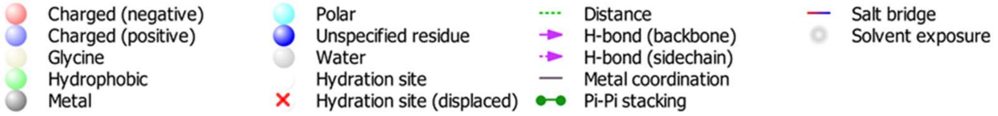

Charged (negative)
Charged (positive)
Glycine
Hydrophobic
Metal
Polar
Unspecified residue
Water
Hydration site
Hydration site (displaced)
Distance
H-bond (backbone)
H-bond (sidechain)
Metal coordination
Pi-Pi stacking
Salt bridge
Solvent exposure

E)

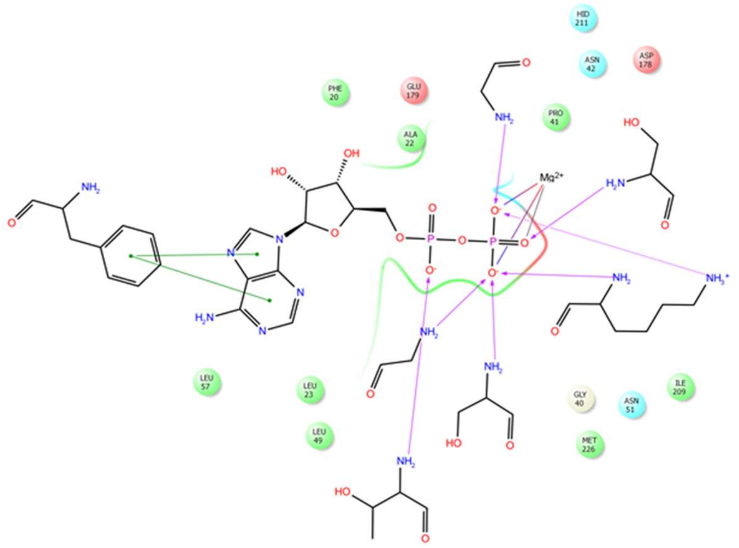
HID 211
ASN 42
ASP 178
PHE 20
GLU 179
PRO 41
ALA 22
Mg2+
LEU 57
LEU 23
LEU 49
GLY 40
ASN 51
ILE 209
MET 226

F)

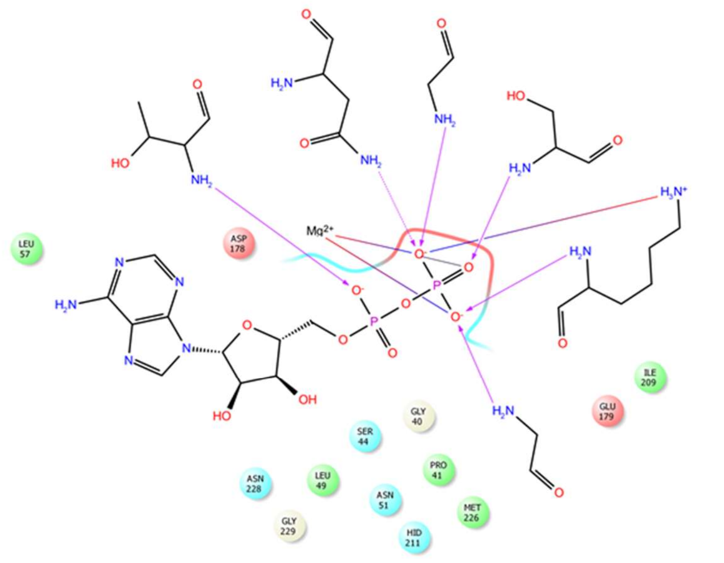
LEU 57
ASP 178
Mg2+
ILE 209
GLU 179
GLY 40
SER 44
PRO 41
ASN 228
LEU 49
ASN 51
MET 226
GLY 229
HID 211

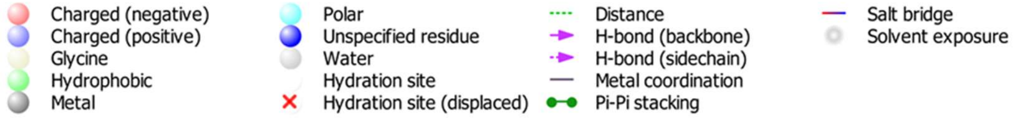
Charged (negative)
Charged (positive)
Glycine
Hydrophobic
Metal
Polar
Unspecified residue
Water
Hydration site
Hydration site (displaced)
Distance
H-bond (backbone)
H-bond (sidechain)
Metal coordination
Pi-Pi stacking
Salt bridge
Solvent exposure

G)

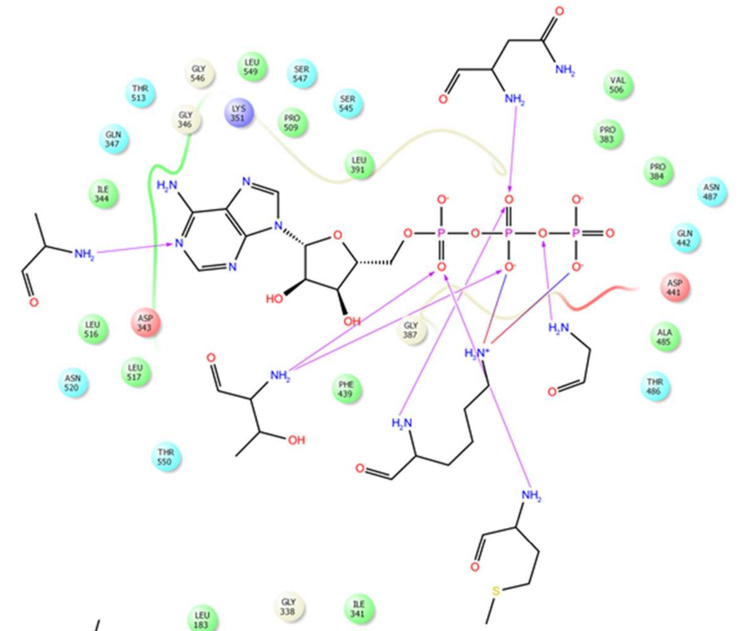

H)

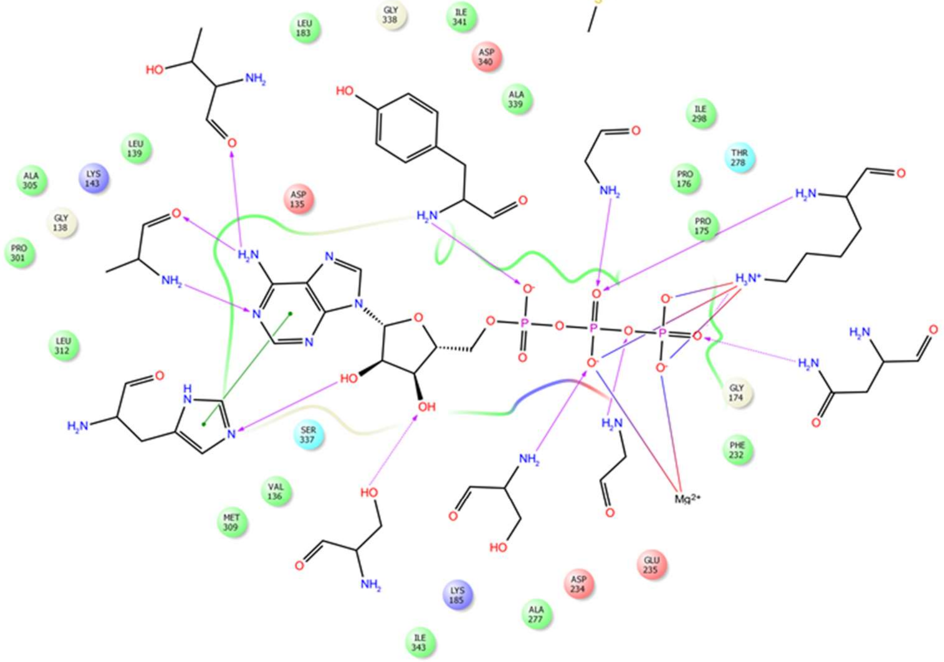

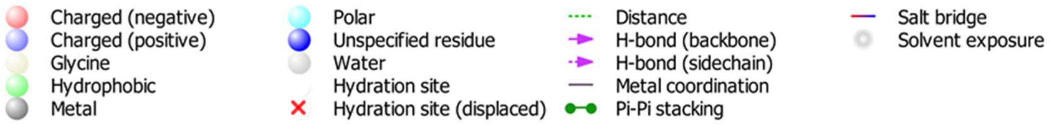
Charged (negative)
Charged (positive)
Glycine
Hydrophobic
Metal
Polar
Unspecified residue
Water
Hydration site
Hydration site (displaced)
Distance
H-bond (backbone)
H-bond (sidechain)
Metal coordination
Pi-Pi stacking
Salt bridge
Solvent exposure

**Fig. S2. The active site analysis of the FlaI protein and its homologues.**

The two dimensional representation of the positioning and interactions between the substrate, cofactor, and amino acids in the active sites of the FlaI protein and its homologues. The selected structures were crystalised with either substrate, substrate analogs or products in the active site. The crystal structure of the FlaI homologues can be found under PDB codes: 5C18 (A), 2QZ4 (B), 4IHQ (C), 4YMV (D), 1G6H (E), 1G9X (F), 5Z6R (G), 2ZAN (H). The interacting amino acids are represented with a chemical structure, and the neighboring amino acids are colour coded based on their influence on the environment.

A.1

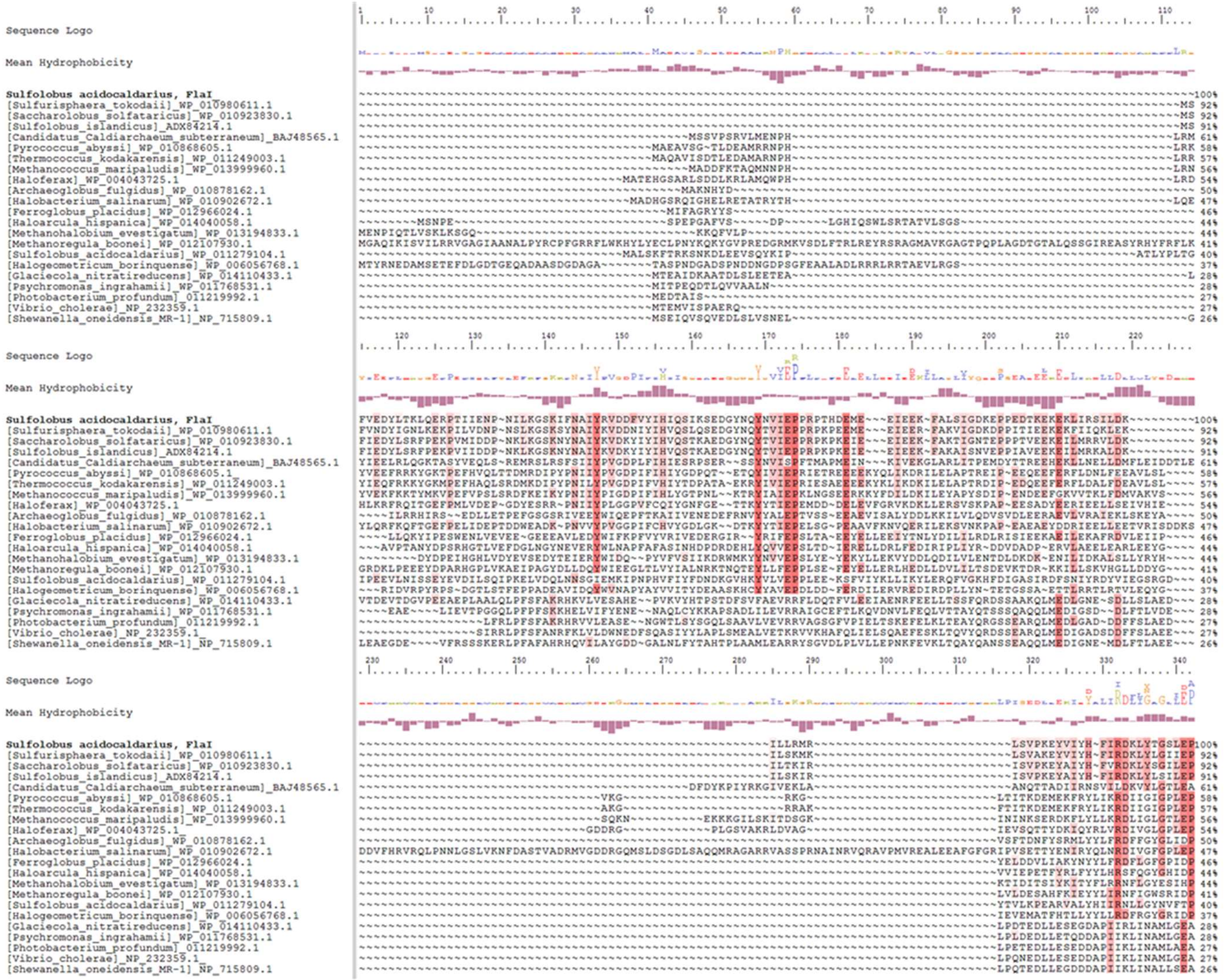
Sequence Logo
Mean Hydrophobicity
Sulfolobus acidocaldarius, FlaI
[Sulfurisphaera_tokodaii]_WP_010980611.1
[Saccharolobus_solfataricus]_WP_010923830.1
[Sulfolobus_islandicus]_ADX84214.1
[Candidatus_Caldiarchaeum_subterraneum]_BAJ48565.1
[Pyrococcus_abyssi]_WP_010868605.1
[Thermococcus_kodakarensis]_WP_011249003.1
[Methanococcus_maripaludis]_WP_013999960.1
[Haloferax]_WP_004043725.1
[Archaeoglobus_fulgidus]_WP_010878162.1
[Halobacterium_salinarum]_WP_010902672.1
[Ferroglobus_placidus]_WP_012966024.1
[Haloarcula_hispanica]_WP_014040058.1
[Methanohalobium_evestigatum]_WP_013194833.1
[Methanoregula_boonei]_WP_012107930.1
[Sulfolobus_acidocaldarius]_WP_011279104.1
[Halogeometricum_borinquense]_WP_006056768.1
[Glaciecola_nitratireducens]_WP_014110433.1
[Psychromonas_ingrahamii]_WP_011768531.1
[Photobacterium_profundum]_011219992.1
[Vibrio_cholerae]_NP_232359.1
[Shewanella_oneidensis_MR-1]_NP_715809.1

A.2

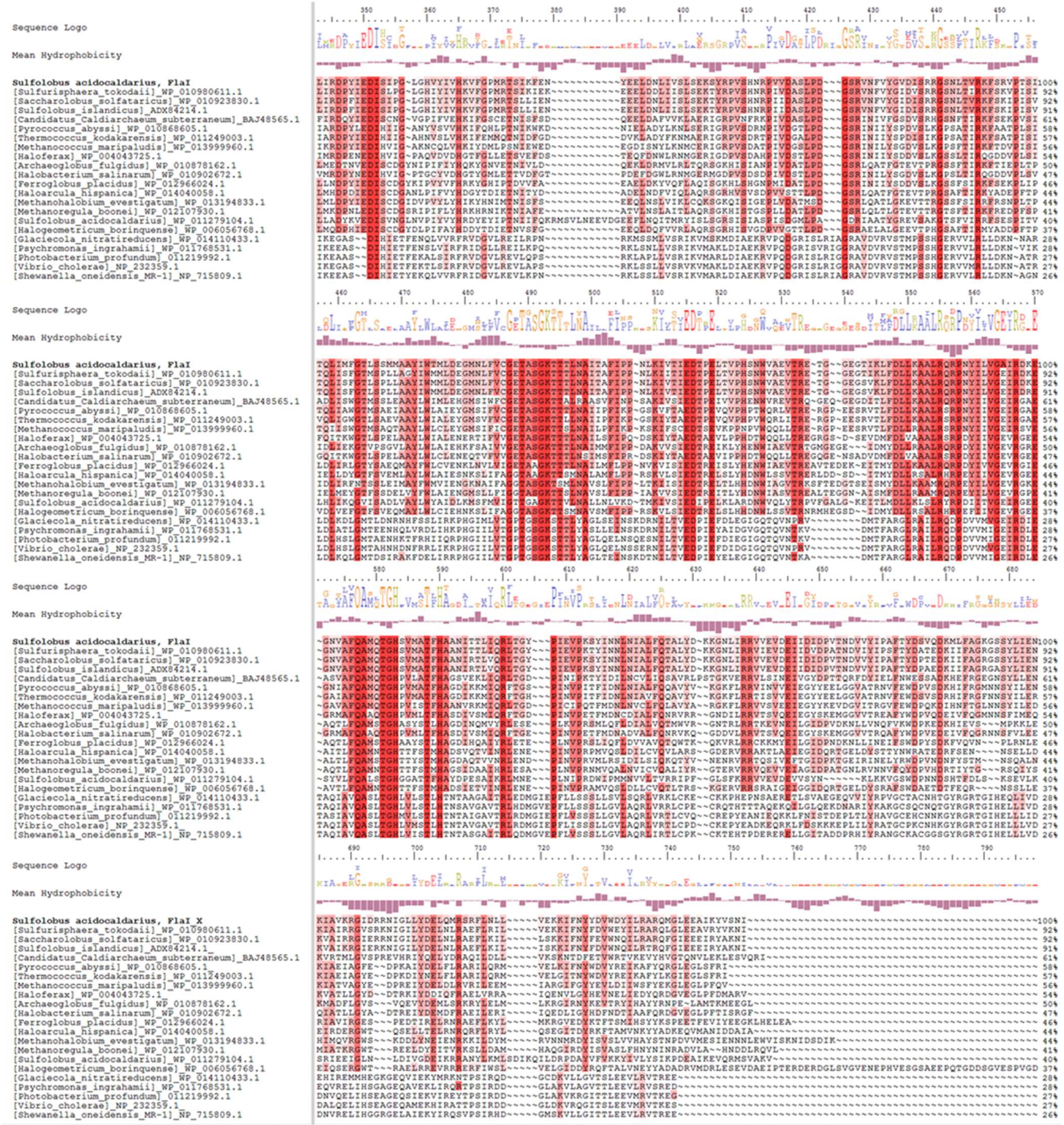
Sequence Logo
Mean Hydrophobicity
Sulfolobus acidocaldarius, FlaI
[Sulfurisphaera_tokodaii]_WP_010980611.1
[Saccharolobus_solfataricus]_WP_010923830.1
[Sulfolobus_islandicus]_ADX84214.1
[Candidatus_Caldiarchaeum_subterraneum]_BAJ48565.1
[Pyrococcus_abyssi]_WP_010868605.1
[Thermococcus_kodakarensis]_WP_011249003.1
[Methanococcus_maripaludis]_WP_013999960.1
[Haloferax]_WP_004043725.1
[Archaeoglobus_fulgidus]_WP_010878162.1
[Halobacterium_salinarum]_WP_010902672.1
[Ferroglobus_placidus]_WP_012966024.1
[Haloarcula_hispanica]_WP_014040058.1
[Methanohalobium_evestigatum]_WP_013194833.1
[Methanoregula_boonei]_WP_012107930.1
[Sulfolobus_acidocaldarius]_WP_011279104.1
[Halogeometricum_borinquense]_WP_006056768.1
[Glaciecola_nitratireducens]_WP_014110433.1
[Psychromonas_ingrahamii]_WP_011768531.1
[Photobacterium_profundum]_011219992.1
[Vibrio_cholerae]_NP_232359.1
[Shewanella_oneidensis_MR-1]_NP_715809.1
Sulfolobus acidocaldarius, FlaI_X

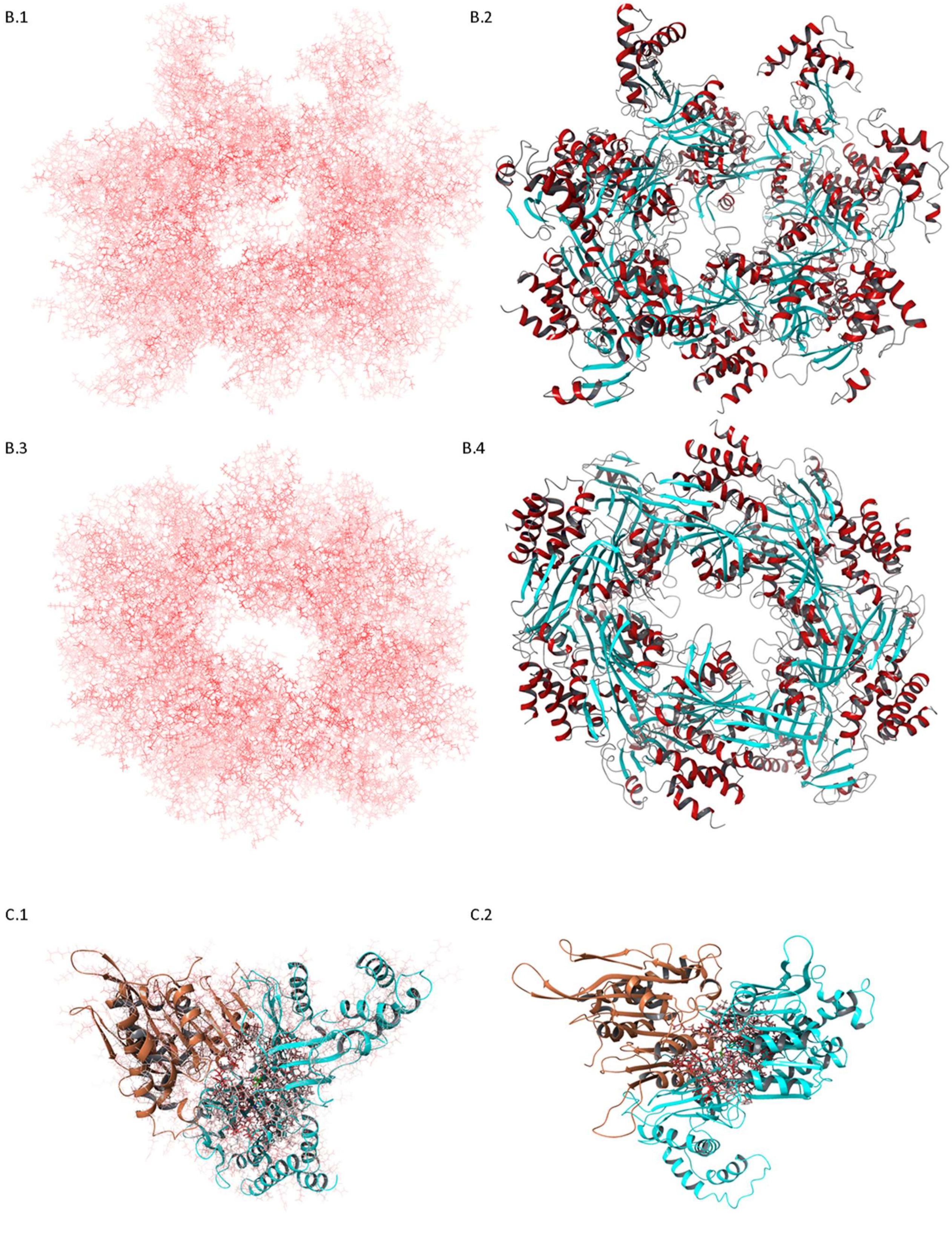
B.1
B.2
B.3
B.4
C.1
C.2

**Fig. S3. The protein sequence alignment of the FlaI family representatives.**

Selection of the key residues and catalytic motifs conserved in the active site of the FlaI family; A. 1-2) The protein sequence alignment of 22 members of FlaI family, with alignment sequence logo and mean hydrophobicity used for recognition of the conserved motifs and key amino acids in the FlaI enzyme. Sequence alignment is color coded by an white-red gradient representing the conservation level of each residue; B) Residue similarity score, in white-red gradient mapped on the structure obtained in molecular dynamics simulations of FlaI (PDB: 4IHQ). Protein complex shown from top (B.1-2) and bottom view (B.3-4). Representations B.2 and B.4 show protein complex with secondary structural elements. Alpha helices and Β-sheets are coloured in red and blue, respectively; C) Structure of the three FlaI domains involved in one active site. Domains belong to two neighboring FlaI monomers coloured in blue and orange, respectively. Represented in licorice model are all amino acids (C.1) and amino acids in the FlaI active site used in further steps (C.2). White-red colour gradient is used to depict evolutionary conservation of each residue of the enzyme.

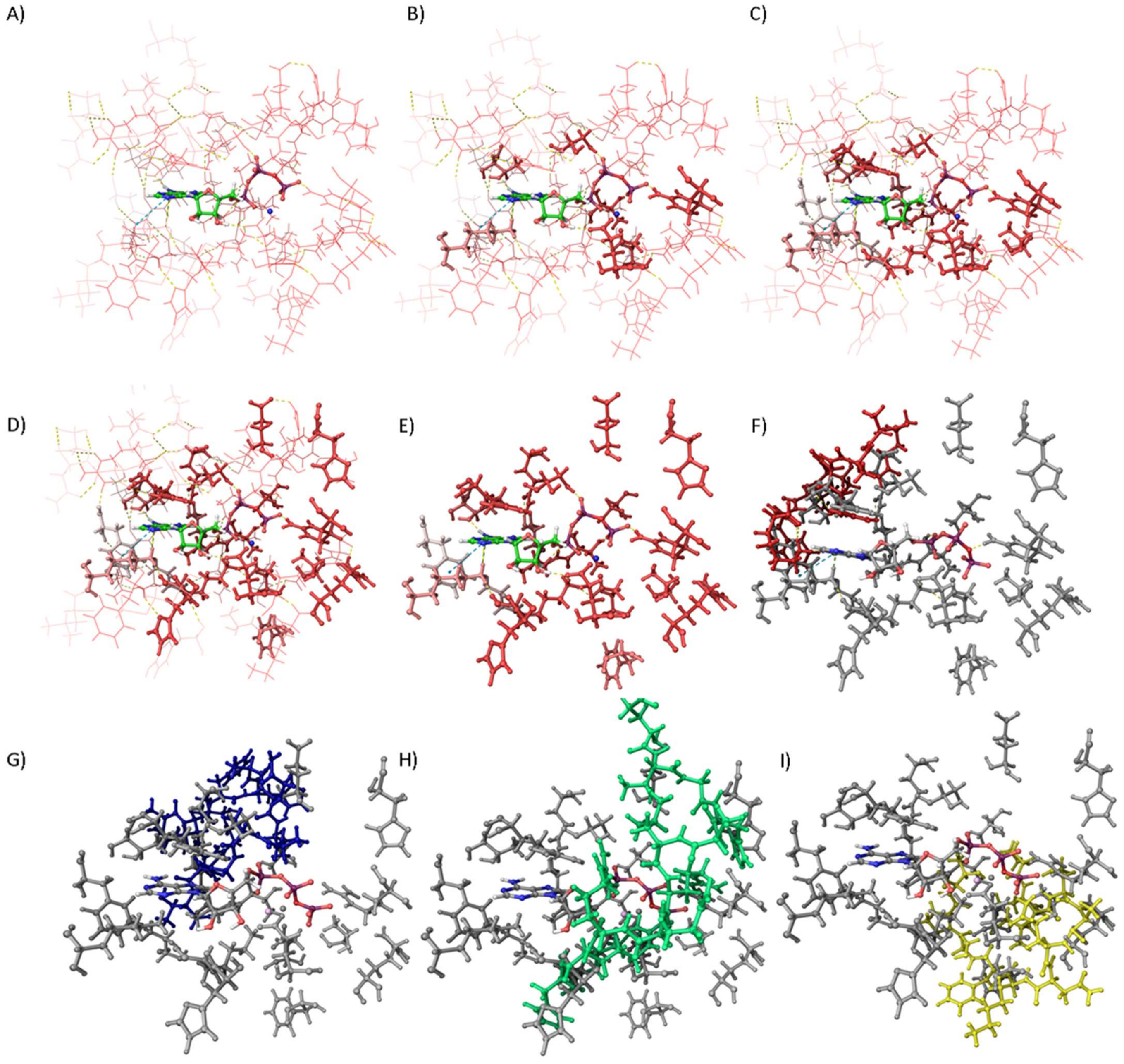
A)
B)
C)
D)
E)
F)
G)
H)
I)

J)

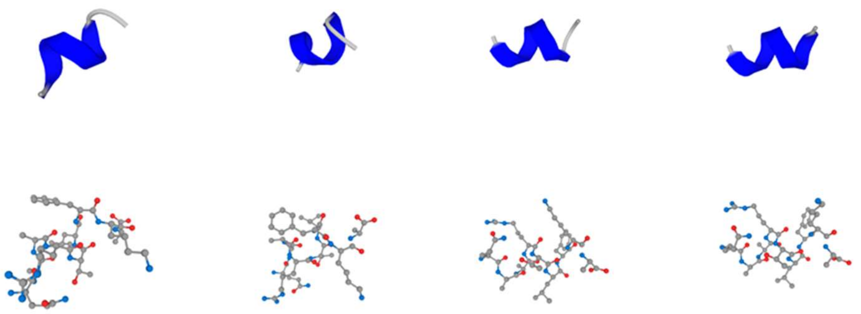

K)

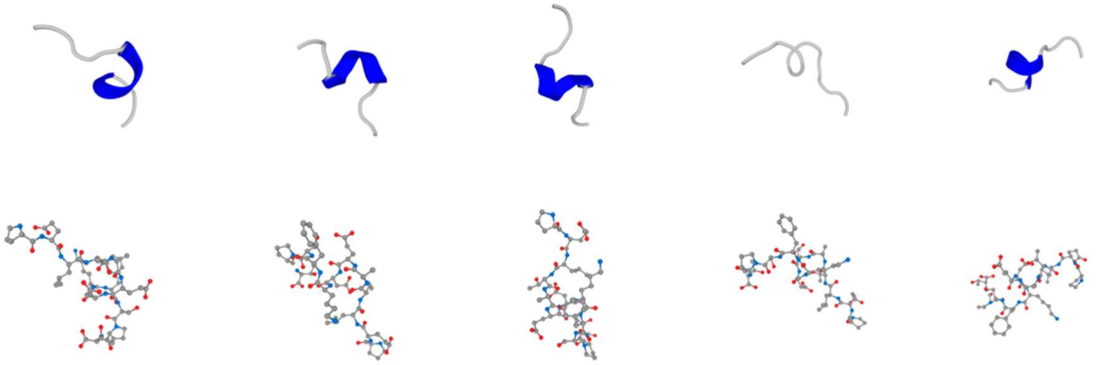

L)

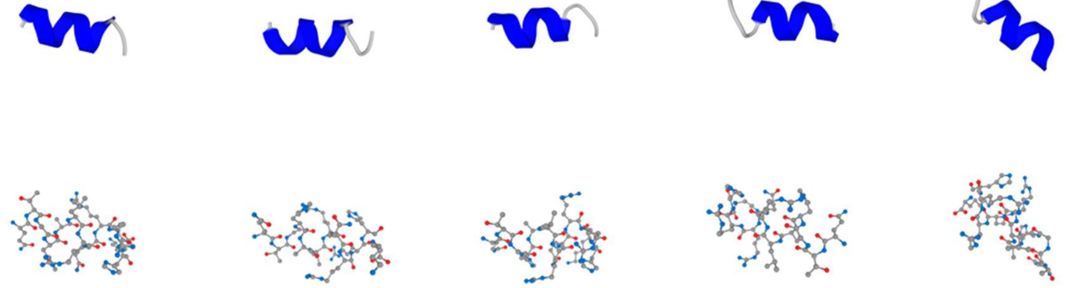

M)

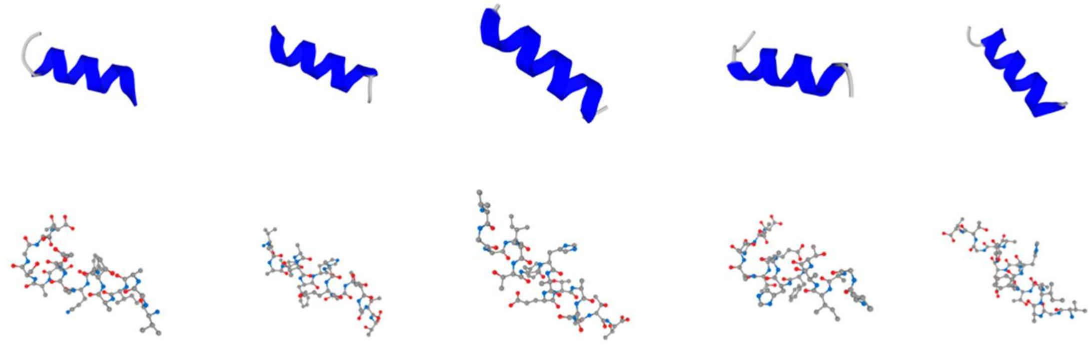

**Fig. S4. Modelling of the active site of the DNA Scaffold-Embedded Protein Emulation Complex (D-SEPEC).**

Through the process of addition of neighboring residues, the key amino acids of the active site of the FlaI enzyme were merged into the four peptides, 10 to 15 amino acids in length; A-E) Stages in the expansion of the emulated active site; F-I) The resulting peptides, served as a base for development of the four peptide-oligonucleotide conjugates (POC 1-4) used in the emulated active center. The peptides are depicted with licorice structure in designated colour: red, yellow, green and blue, respectively. J-M) Cluster representatives of the folding simulations were used as the most occurring conformations of each peptide element. The most occurring conformations of peptide elements of the emulated catalytic site obtained from folding simulations. Each conformation is represented by the atomic and cartoon model. Peptide folding prediction was done by using the PEP-FOLD3 [44].

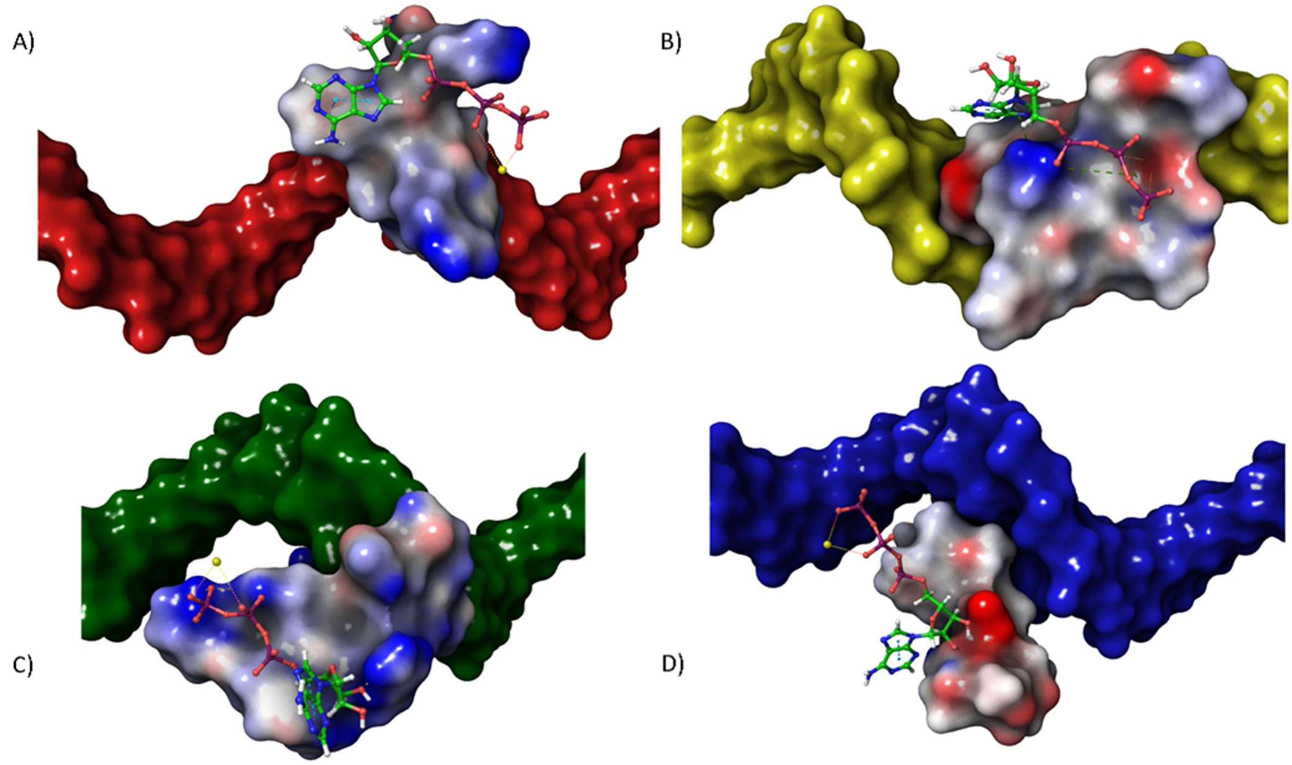

**Fig. S5. Interactions between substrate and peptide-oligonucleotide conjugates (POC).**

The peptide elements are linked with the oligonucletoide elements into the peptide oligonucleotide conjugates (POC 1 - 4); A-D) The POCs interacts with specific areas of the substrate or cofactor. Peptide elements are represented by the VDW surface coloured in the partial charges. The oligonucleotide elements, represented by the VDW surface in designated colors for each POC. The oligonucleotide elements retained their ds DNA conformation by hybridisation with the short Carrier Strand fragments (not shown). Structure of substrate and cofactor is represented by licorice model.

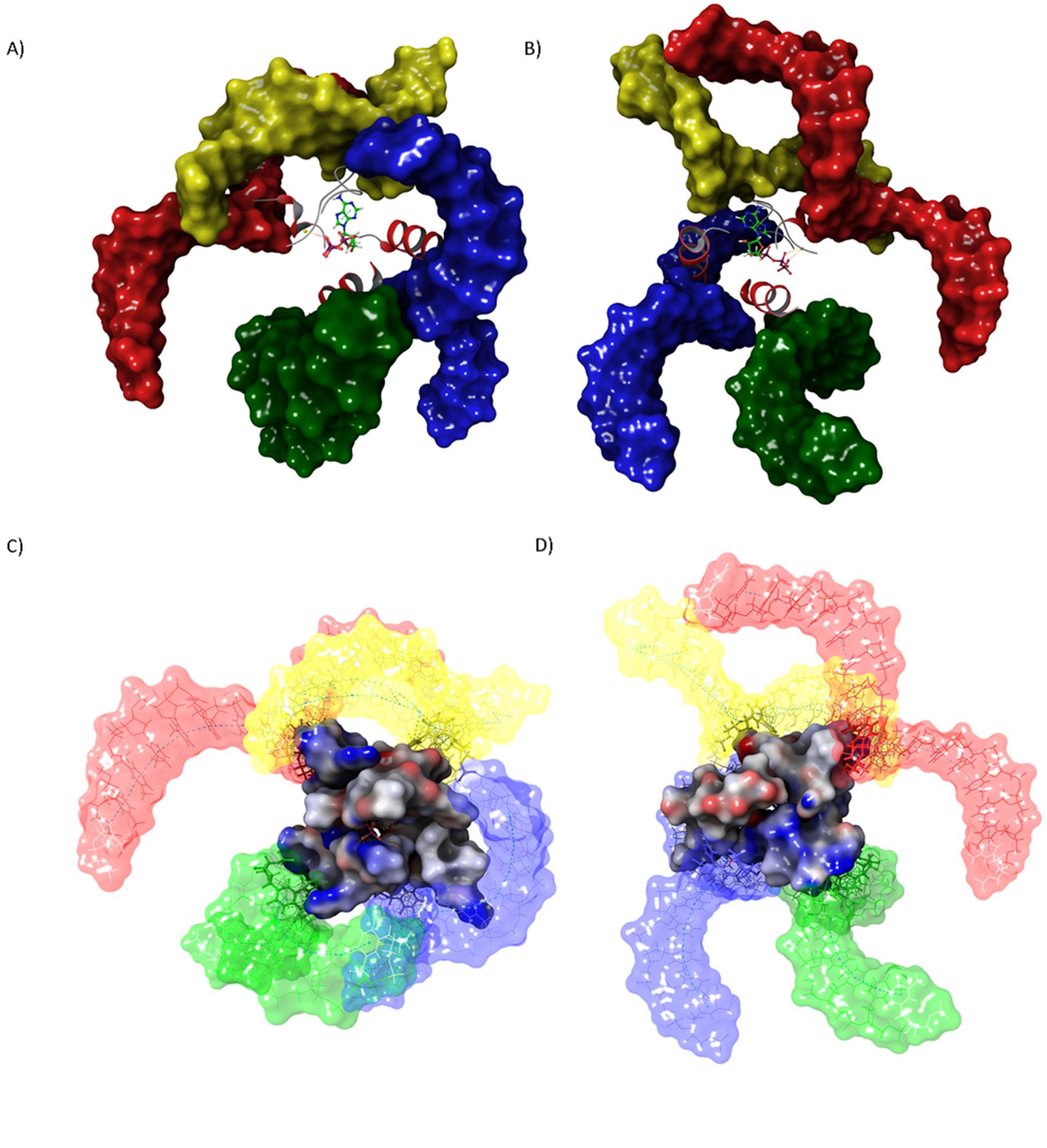
A)
B)
C)
D)

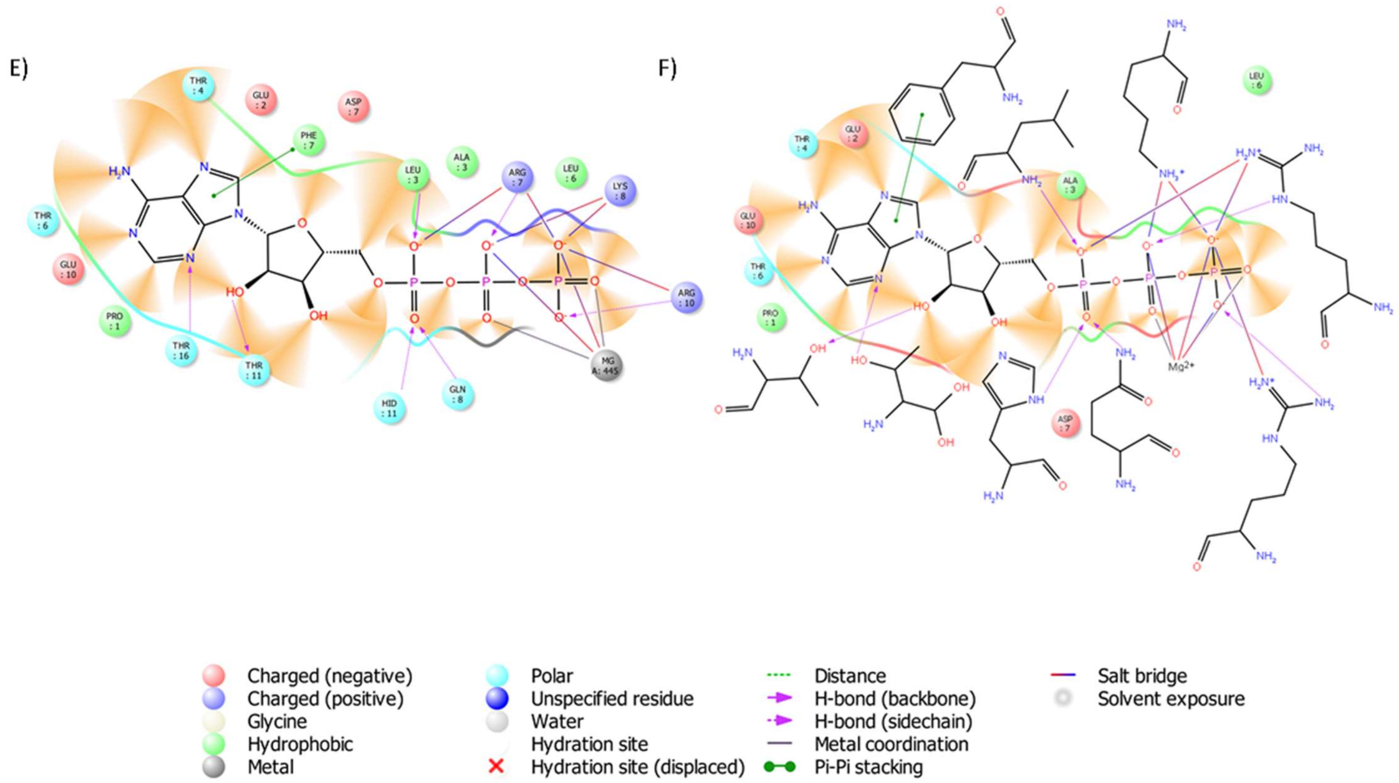

**Fig S6. The assembly of the emulated active site of DNA Scaffold-Embedded Protein Emulation Complex (D-SEPEC).**

The POC elements are combined to form an emulated active site; A-D) Following the optimisation process, this model was used for *in silico* integration on the Carrier Strands of the DNA-origami nanostructure. Peptide elements are represented by the cartoon model (A,B), and VDW surface coloured in the partial charges (C,D). The oligonucleotide elements, represented by the VDW surface in designated colors for each POC; E-F) Two-dimensional representation of the positioning and interactions between the substrate, cofactor, and amino acids in the emulated active site of the D-SEPEC nanozyme. Interacting amino acids are represented by the type (E) or chemical structure (F), while neighboring amino acids are colour coded based on their influence on the environment.

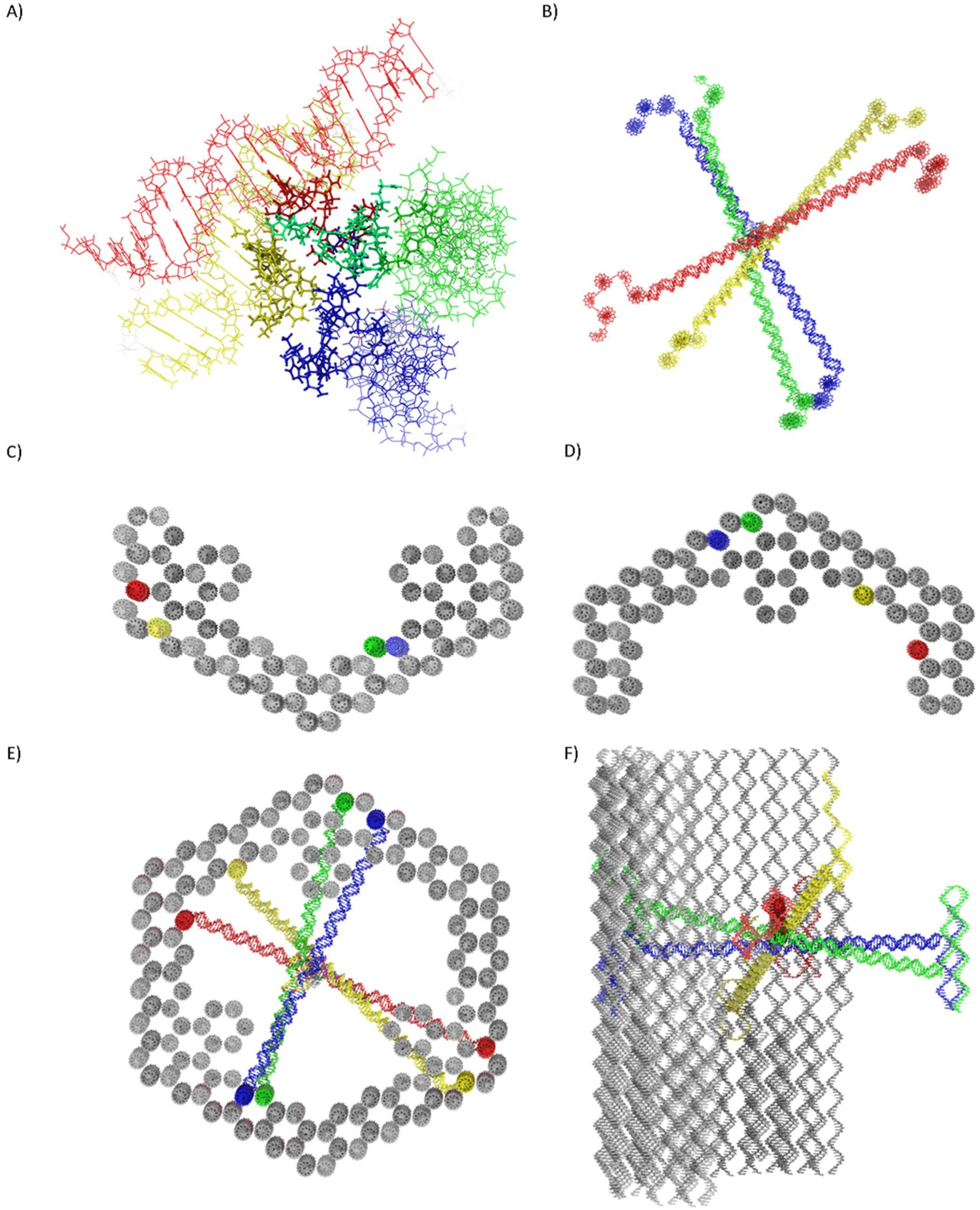
A)
B)
C)
D)
E)
F)

**Fig. S7. The super assembly of the DNA Scaffold-Embedded Protein Emulation Complex (D-SEPEC).**

Main structural elements required for the super assembly of the D-SEPEC nanozyme. A) Each POC element of the emulated catalytic site, in predefined relative position and orientation, was hybridised with a 20 bp long, Carrier Strand fragment, represented by the all-atom model of dedicated color; B) Carrier Strands were modelled by the linear prolongation of the Carrier Strand fragments. Once prolonged, Carrier Strands were adjusted relative to the postion of the Shell structure in a way that both ends of each Carrier Strand overlap with the staple strands protruding from the inner surface area of the Shell structure. Carrier Strands were comprised from five to six ssDNA elements, each holding two or more domains complementary to other elements of the structure (see tab. 4). Once the optimal Shell's staple strands were chosen, a caDNAno software was used to extract the sequences and adjust the surounding staple strands in the Shell structure (tab. 4). Staple strands were merged with the appropriate elements of the Carrier Strands (tab. 5); C-D) Elements of the hollow, hexagonal, Shell structures, Sp8064 and Sp7569, were designed separately in an iterative procedure using the caDNAno and CanDo program packages [35,45]. To form a single nanostructure, Shell elements were designed on the principles of shape complementarity and bridging strands. All-atom model of the Shell structures was created using the cadnano to PDB converter [46]. Folding and sequence fragmentation artefacts, caused by the data transformation, were corrected manually and stabilised by the gradient descent energy minimisation cycles using the SCHRODINGER program package[42]. Carrier Strand integration areas are represented by the red, yellow, green and blue color respectively; E) Super assembly of the D-SEPEC nanozyme; F) Side view of the D-SEPEC nanozyme lacking the SP7560 Shell element.

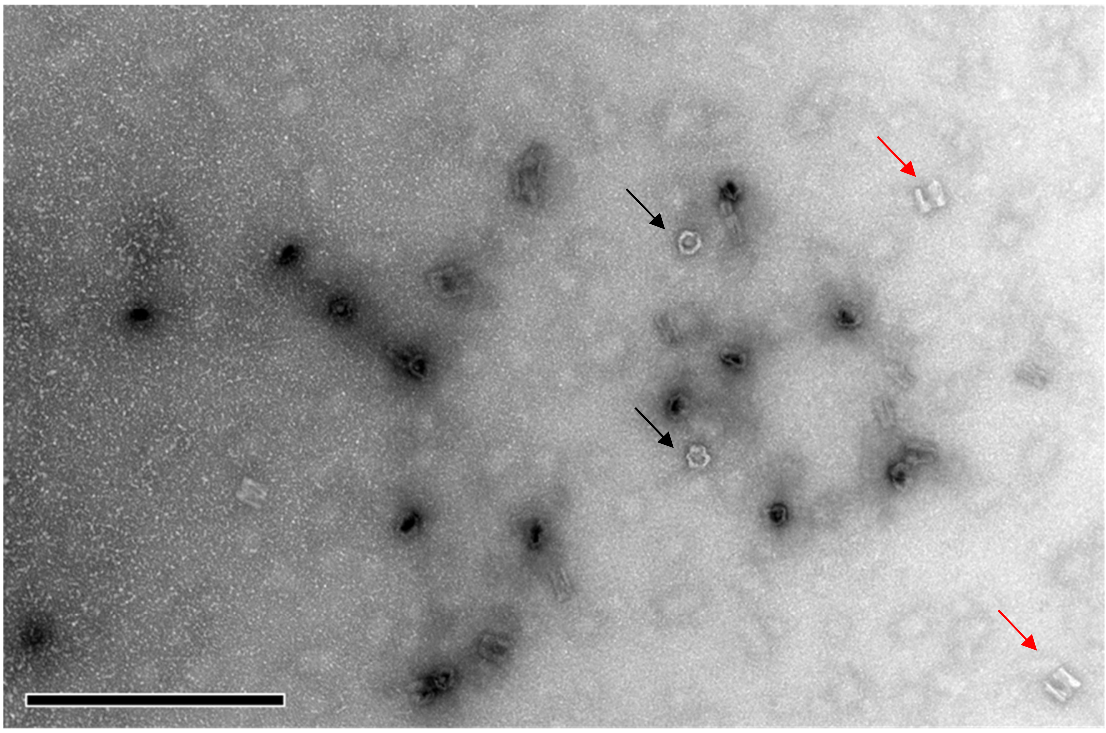

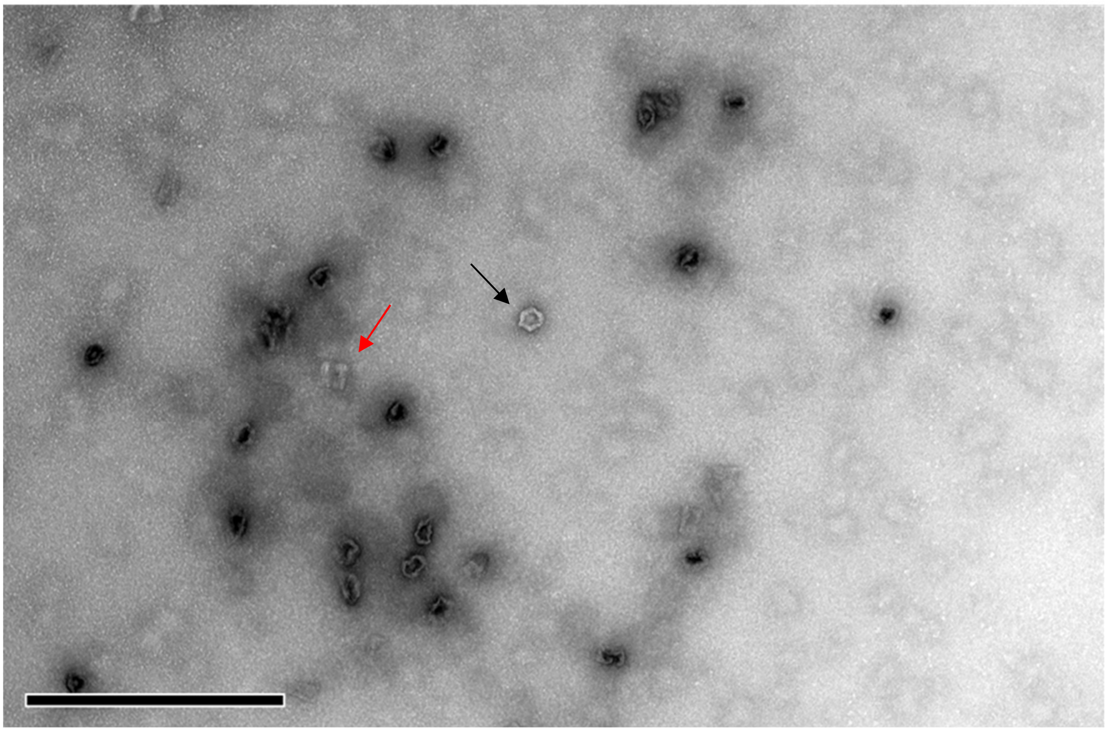

**Fig. S8. The negative staining transmission electron microscopy (nsTEM).**

The super assembly of the Shell structure was confirmed by the nsTEM imaging. Stability of the structure is visible by its correlations with the predefined dimensions and shape both in top view (black arrow) and side-view (red arrow) of major structural elements. Due to the process of TEM imaging the majority of observed constructs show structural deformations. Scale bar, 500 nm.

A)

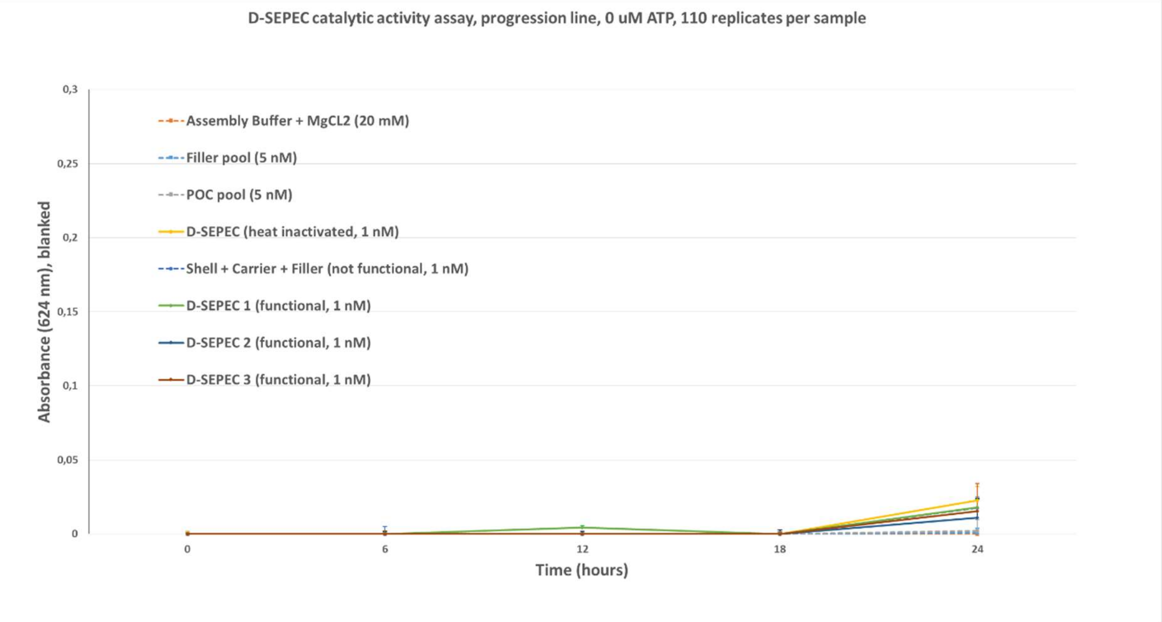
D-SEPEC catalytic activity assay, progression line, 0 uM ATP, 110 replicates per sample
Assembly Buffer + MgCL2 (20 mM)
Filler pool (5 nM)
POC pool (5 nM)
D-SEPEC (heat inactivated, 1 nM)
Shell + Carrier + Filler (not functional, 1 nM)
D-SEPEC 1 (functional, 1 nM)
D-SEPEC 2 (functional, 1 nM)
D-SEPEC 3 (functional, 1 nM)
Absorbance (624 nm), blanked
0,3
0,25
0,2
0,15
0,1
0,05
0
0
6
12
18
24
Time (hours)

B)

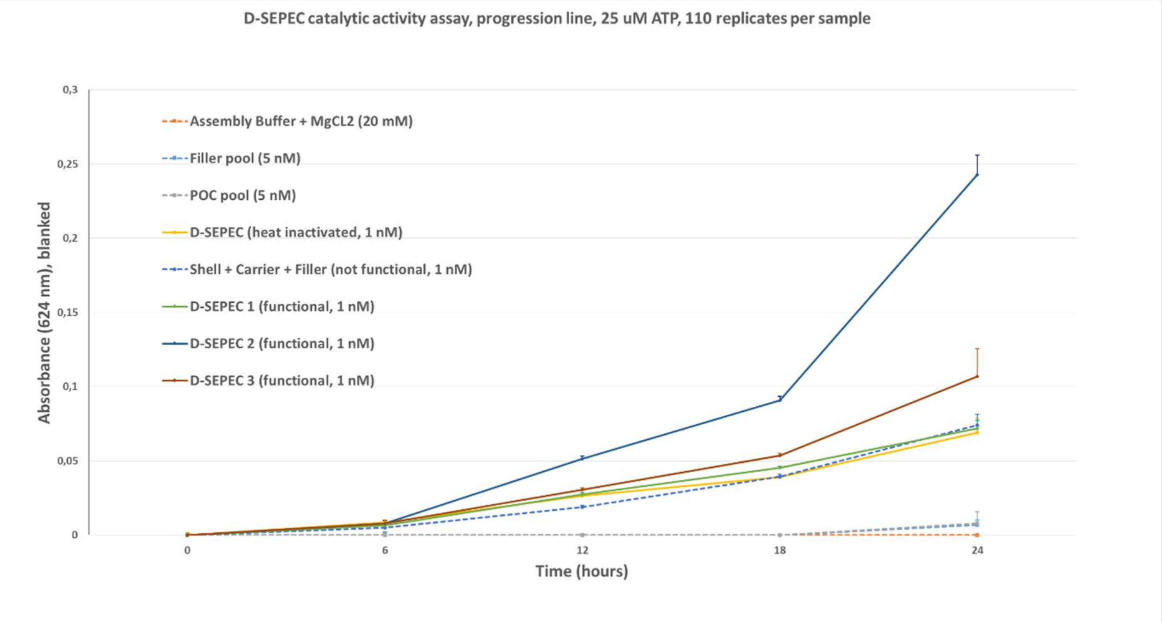
D-SEPEC catalytic activity assay, progression line, 25 uM ATP, 110 replicates per sample
Assembly Buffer + MgCL2 (20 mM)
Filler pool (5 nM)
POC pool (5 nM)
D-SEPEC (heat inactivated, 1 nM)
Shell + Carrier + Filler (not functional, 1 nM)
D-SEPEC 1 (functional, 1 nM)
D-SEPEC 2 (functional, 1 nM)
D-SEPEC 3 (functional, 1 nM)
Absorbance (624 nm), blanked
0,3
0,25
0,2
0,15
0,1
0,05
0
0
6
12
18
24
Time (hours)

C)

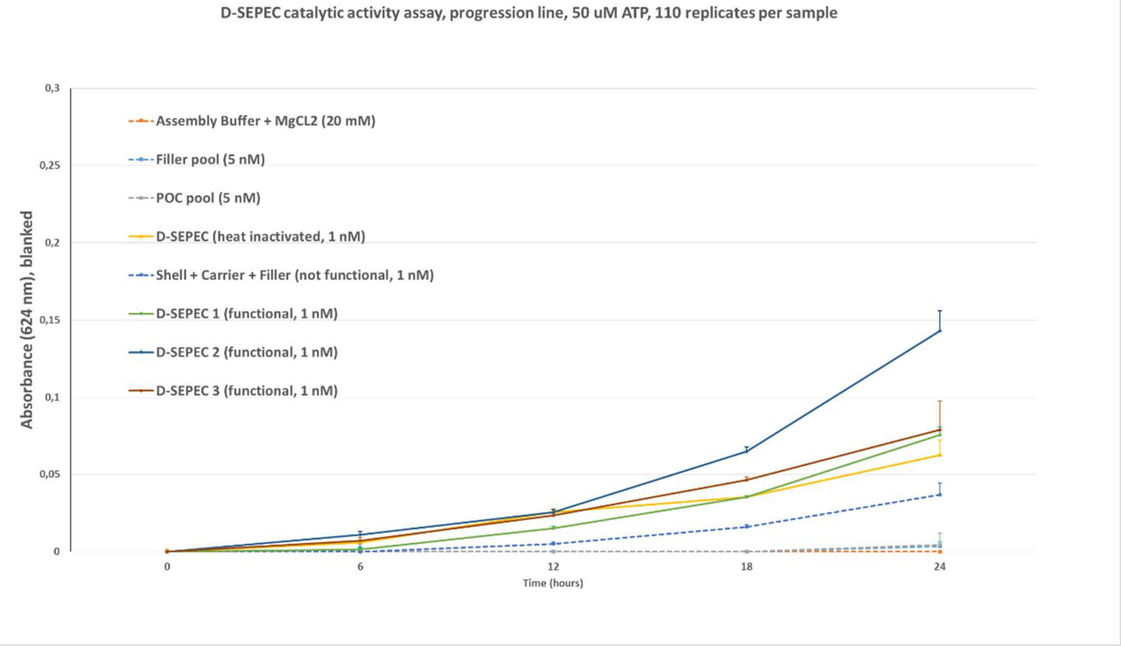

**Fig. S9. The catalytic activity measurements**

The catalytic activity was measured in discrete time intervals in reactions with 0 (A), 25 µM (B) and 50 µM (C) ATP using the Malachite Green assay. The auto hydrolysis of ATP in the assembly buffer was used to blank the values. The samples with the D-SEPEC nanozyme exhibited the strongest catalytic activity (kcat values of 0.2271 s-1 in systems incubated with 25 µM of ATP, and 0.3622 s-1 in systems with 50 µM of ATP). Heat inactivation of the functional D-SEPEC nanodevice (yellow) is marked by a significant activity drop, close to the levels observed with the non-functional DNA nanostructure (Shell, Carrier and Filler; blue, dotted).

A)

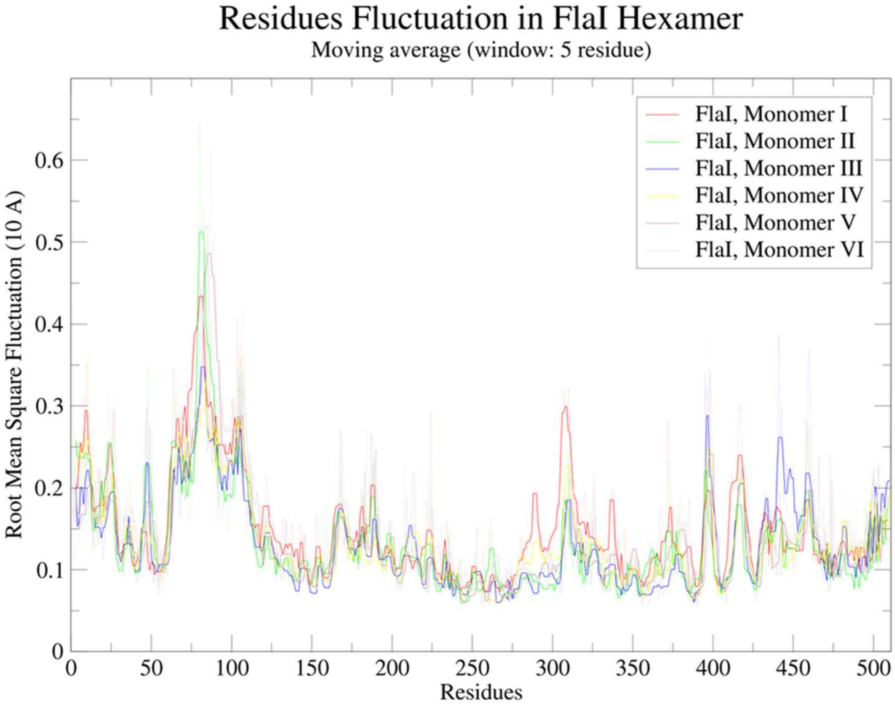
Residues Fluctuation in FlaI Hexamer
Moving average (window: 5 residue)
Root Mean Square Fluctuation (10 A)
Residues
FlaI, Monomer I
FlaI, Monomer II
FlaI, Monomer III
FlaI, Monomer IV
FlaI, Monomer V
FlaI, Monomer VI
0
0.1
0.2
0.3
0.4
0.5
0.6
50
100
150
200
250
300
350
400
450
500

B)

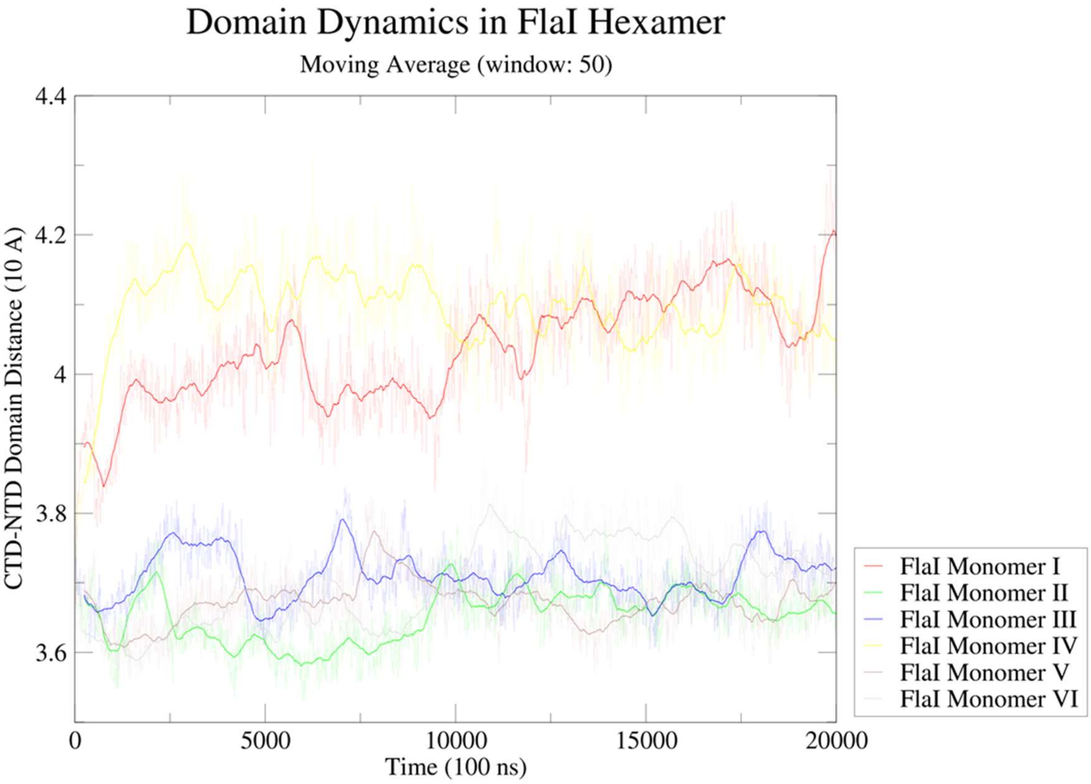
Domain Dynamics in FlaI Hexamer
Moving Average (window: 50)
CTD-NTD Domain Distance (10 A)
Time (100 ns)
FlaI Monomer I
FlaI Monomer II
FlaI Monomer III
FlaI Monomer IV
FlaI Monomer V
FlaI Monomer VI
3.6
3.8
4
4.2
4.4
0
5000
10000
15000
20000

c)

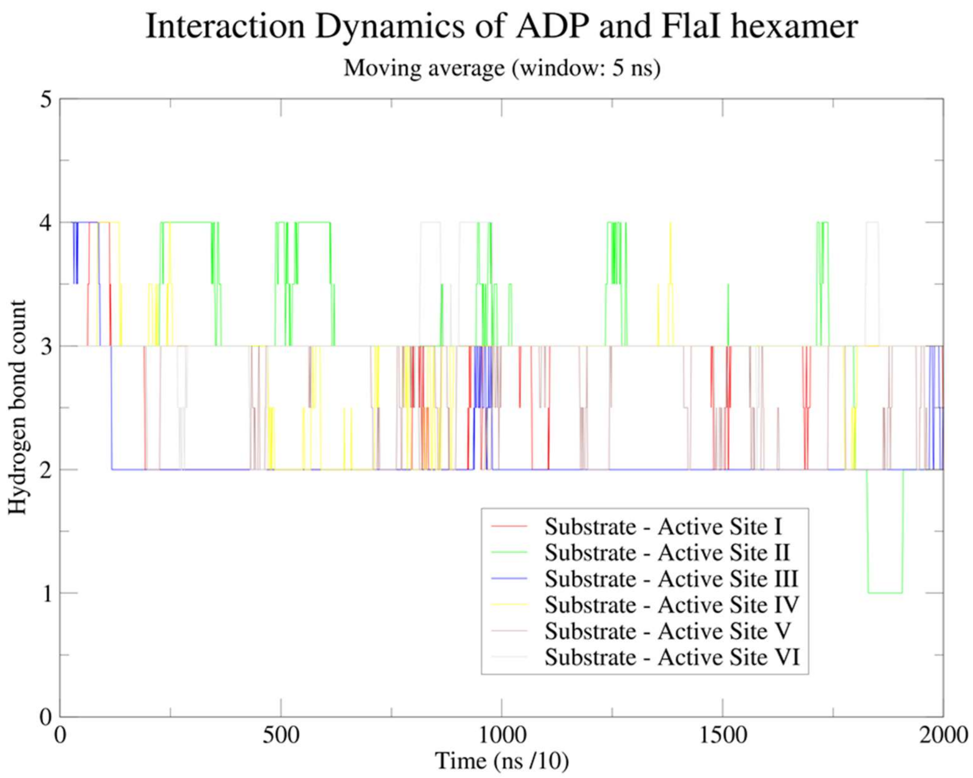

**Fig. S10. The molecular dynamics of the FlaI protein complex observed in the simulations.** Dynamics of residue (A), domains (B) and non-covalent interactions (C) observed through 200 ns of the MD simulations of the FlaI hexamer; A) Comparison of the residue's fluctuation between the six monomers of the FlaI hexamer; B) Dynamics of the FlaI monomers shown as a change in distances between centers of mass of CTD and NTD domains; C) Comparison of the hydrogen bonds forming dynamics between the substrate and active sites of the FlaI hexamer.

| | |
|---|---|
| **POC 1** | Gln**Ala***AlaArgThrLeuPheLys |
| | GCATAGGCTG**T***CGAGGGTCG |
| **POC 2** | ProGluLeuThr**Ala***GluAspLysPheSerProThr |
| | CCCAGACCGC**T***GAAGATCGG |
| **POC 3** | GlnThrGlyHis**Ala***LeuArgGlnArgArgHisLys |
| | CCATCAGGG**T***GCGGTCTGGC |
| **POC 4** | ValGlyAlaIleAlaThrPheHis**Ala***GluThrAlaSerGlyThrThr |
| | CGGGTCGGTCAGG**T***GGTAATG |

**Table S1. Sequences of the peptide oligonucleotide conjugates used in this study.**

Linker positions are marked by Ala* (L-azidohomoalanine residue) in peptide elements, and T* (5-ethynyl-2'deoxyuridine) in oligonucleotide elements.

| **POC 1** | Gln | Ala* | Ala | Arg | Thr | Leu | Phe | Lys |
|---|---|---|---|---|---|---|---|---|
| Native | Gln | Ala | Ala | Arg | Thr | Gly | Phe | Lys |
| Alignment # | 630 | | | 643 | 469 | 468 | 467 | 371 |
| Res.ID # | 399 | | | 412 | 238 | 237 | 236 | 140 |

| **POC 2** | Pro | Glu | Leu | Thr | Ala* | Glu | Asp | Lys | Phe | Ser | Pro | Thr |
|---|---|---|---|---|---|---|---|---|---|---|---|---|
| Native | Pro | Glu | Leu | Thr | Ala | Glu | Asp | Lys | Phe | Ser | Leu | Thr |
| Alignment # | 523 | 524 | 525 | 518 | 519 | 520 | 521 | 453 | 454 | 455 | 456 | 457 |
| Res.ID # | 292 | 293 | 294 | 287 | 288 | 289 | 290 | 223 | 224 | 225 | 226 | 227 |

| **POC 3** | Gln | Thr | Gly | His | Ala* | Leu | Arg | Gln | Arg | Arg | His | Lys |
|---|---|---|---|---|---|---|---|---|---|---|---|---|
| Native | Gln | Thr | Gly | His | Ala | Leu | Arg | Gln | Arg | Arg | His | Lys |
| Alignment # | 585 | 586 | 587 | 588 | 558 | 559 | 560 | 561 | 562 | | | |
| Res.ID # | 350 | 351 | 352 | 353 | 324 | 325 | 326 | 327 | 328 | | | |

| **POC 4** | Val | Gly | Ala | Ile | Ala | Thr | Phe | His | Ala* | Glu | Thr | Ala | Ser | Gly | Thr | Thr |
|---|---|---|---|---|---|---|---|---|---|---|---|---|---|---|---|---|
| Native | Val | | | Leu | Ala | Thr | Phe | His | Ala | Glu | Thr | Ala | Ser | Gly | Thr | Thr |
| Alignment # | 585 | | | 586 | 587 | 588 | 589 | 590 | 591 | 488 | 489 | 490 | 491 | 492 | 493 | 494 |
| Res.ID # | 591 | | | 592 | 593 | 594 | 595 | 596 | 597 | 263 | 264 | 265 | 266 | 267 | 269 | 270 |

**Table S2. The retained residues of the native FlaI protein complex.**

Residues of the native *Sulfolobus acidocaldarius* FlaI, obtained after 200 ns of MD simulation that were used in the construction of the POC oligonucleotides. Res.ID numbers belong to the residues of the crystal structure (PDB: 4iHQ) [3]. Following the design process some residues were changed or could not be aligned with the native structure, and are represented by the blank cells.

| | |
|---|---|
| **POC 1 Filler** | GCATAGGCTGTCGAGGGTCG |
| **POC 2 Filler** | CCCAGACCGCTGAAGATCGG |
| **POC 3 Filler** | CCATCATGGGTGCGGTCTGGC |
| **POC 4 Filler** | CGGGTCGGTCAGGTGGTAATG |

**Table S3. Sequences of the Filler molecules used as the non –functional variants of the POCs.**

To create the non-functional D-SEPEC nanozymes, in the process of super assembly, the POC molecules were replaced by four oligonucleotides with the identical sequence.

| | |
|---|---|
| **POC 1 A** | CAATAATCGGCTGTTTAGCGAACCTCCATCTTACCAACGCTAACGCATGTGGAATTAGAGGCC |
| **POC 1 B** | ATGTGACGAGCTTCATTTATACGCTTCGCGCCGACCCTCGACAGCCTATGCGCGCAGCAGTGCACAAG |
| **POC 1 C** | CGTACGATGCAACTGTGTGG AGCTTCAAAGACCTTCATTTAAACAGTTCAGAGAATCCCCCTCAAA |
| **POC 1 D** | GCGCGAAGCGTATAAATGAAGCTCGTCACATGGCCTCTAATTCCACATGCG |
| **POC 1 E** | CCACACAGTTGCATCGTACGCTTGTGCACTGCTGCGC |
| **POC 2 A** | CCCAATCCAAAGTCAGCCCAATTCGCCTAATTATTACCGCATCTTGACCTAACTG |
| **POC 2 B** | ACAAACTGCCATGGACGACTAGCCATGCCCGATCTTCAGCGGTCTGGGATTGGCTCTTAGACAGCCCGATACAGTGATT |
| **POC 2 C** | GGCGCATCCTGATAAATTGTGAATACAATCACTGTATCGGGCTGTCTAAGAGCCAAT |
| **POC 2 D** | GCATGGCTAGTCGTCCATGGCAGTTTGTCAGTTAGGTCAAGATGCGGT |
| **POC 3 A** | GATAAAGCATTCCGCTCGGGTATGGCAGTAAGTACGCCTTCTGAATTGGCCAGACCGCACCCATGATGGTGCTAACCTTCAT |
| **POC 3 B** | CTCGCTCACATTGAGCTACTGATAGGTCACGTTGGTGGCCATCACAAGGCG |
| **POC 3 C** | CAATTCAGAAGGCGTACTTACTGCCATACCCGAGCGGAATGCTTTATCTTGCGATTTCTTATTAGAGAGATAGA |
| **POC 3 D** | AGTAGCTCAATGTGAGCGAGATGAAGGTTAGCA |
| **POC 4 A** | ACTCATCAAGAGGCAAAGACAGCATACTGCGCTGTATAGTCGAAT |
| **POC 4 B** | CGATTAATACAACGAACGGTGATGTTGTCCATTACCACCTGACCGACCCG ATAGATTCGGATAACATGATTAGCCGAAGT |
| **POC 4 C** | GACAACATCACCGTTCGTTGTATTAATCGATTCGACTATACAGCGCAGT |
| **POC 4 D** | GTCGTGTTAGGGAAATTAGCCTGTGAAGCCGTCTACTTCGGCTAATCATGTTATCCGAATCTAT |

**Table S4. Sequences of the Carrier Strand elements used in this study.**

Carrier Strands are comprised from the several single stranded oligonucleotides designed to form longer, linear ds DNA structure capable of integrating on both ends into the Shell nanostructure by replacing preselected staple strands. Once formed, each Carrier Strand hold a single stranded domain complementary to one POC and Filler pair.

| POC 1 | Shell integration | | | POC / Filler binding site | | | Shell integration |
|---|---|---|---|---|---|---|---|
| 5′ | CAATAATCGGCTGTTTTAGCGAACCTCCATCTTACCAACGCTAA | CGCATGTGGAATTAGAGGCC | ATGTGACGAGCTTCATTTATACGCTTCGCGC | CGACCCTCGACAGCCTATGC | GCGCAGCAGTGCACAAG | CGTACGATGCAACTGTGTGG | AGCTTCAAAGACCTTCATTTAAACAGTTCAGAGAATCCCCCTCAAA |
| 3′ | | GCGTACACCTTAATCTCCGG | TACACTGCTCGAAGTAAATATGCGAAGCGCG | GCTGGGAGCT*GTCGGATACG | CGCGTCGTCACGTGTTC | GCATGCTACGTTGACACACC | |

| POC 2 | Shell integration | | | POC / Filler binding site | | Shell integration |
|---|---|---|---|---|---|---|
| 5′ | CCCAATCCAAAGTCAGCCCAATTCGCCTAATTATT | ACCGCATCTTGACCTAACTG | ACAAACTGCCATGGACGACTAGCCATGC | CCGATCTTCAGCGGTCTGGG | ATTGGCTCTTAGACAGCCCGATACAGTGATT | |
| 3′ | | TGGCGTAGAACTGGATTGAC | TGTTTGACGGTACCTGCTGATCGGTACG | GGCTAGAAGT*CGCCAGACCC | TAACCGAGAATCTGTCGGGCTATGTCACTAA | CATAAGTGTTAAATAGTCCTACGCGG |

| POC 3 | Shell integration | | | POC / Filler binding site | | | Shell integration |
|---|---|---|---|---|---|---|---|
| 5′ | | GATAAAGCATTCCGCTCGGG | TATGGCAGTAAGTACGCCTTCTGAATTG | GCCAGACCGCACCCATGATGG | TGCTAACCTTCAT | CTCGCTCACATTGAGCTACT | GATAGGTCACGTTGGTGGCCATCACAAGGCG |
| 3′ | AGATAGAGAGATTATTCTTTAGCGTT | CTATTTCGTAAGGCGAGCCC | ATACCGTCATTCATGCGGAAGACTTAAC | CGGTCTGGCGT*GGGTACTACC | ACGATTGGAAGTA | GAGCGAGTGTAACTCGATGA | |

| POC 4 | Shell integration | | | POC / Filler binding | | Shell integration |
|---|---|---|---|---|---|---|
| 5′ | ACTCATCAAGAGGCAAAGACAGCAT | ACTGCGCTGTATAGTCGAAT | CGATTAATACAACGAACGGTGATGTTGTC | CATTACCACCTGACCGACCCG | ATAGATTCGGATAACATGATTAGCCGAAGT | |
| 3′ | | TGACGCGACATATCAGCTTA | GCTAATTATGTTGCTTGCCACTACAACAG | GTAATGGT*GGACTGGCTGGGC | TATCTAAGCCTATTGTACTAATCGGCTTCA | TCTGCCGAAGTGTCCGATTAAAGGGATTGTGCTG |

**Table S5. The domain alignment of the Carrier Strand elements.**

Each Carrier Strand hold a Shell integration sequence on both ends. These sequences take place of the staple strand by the complementary interaction with the scaffolds of the Shell structure. In the central position of each Carrier Strand is the POC / Filler binding site which position and orientation directly influence position and orientation of POC elements of the D-SEPEC.

**REFERENCES AND NOTES**

39. M. J. Abraham *et al*., GROMACS: High performance molecular simulations through multi-level parallelism from laptops to supercomputers. SoftwareX 1, 19-25 (2015). doi: 10.1016/j.softx.2015.06.001
D.A. Case *et al.,* AMBER 2016, University of California, San Francisco (2016).
40. Meagher, *et al*., Development of polyphosphate parameters for use with the AMBER force field. J. Comp. Chem. 24, 1016 (2003). doi: 10.1002/jcc.10262
41. O. Allnér, *et al.,* Magnesium Ion-Water Coordination and Exchange in Biomolecular Simulations, J. Chem. Theory Comput. 8, 1493-1502 (2012). doi: 10.1021/ct3000734
42. Schrödinger Release 2019-3: Maestro, Schrödinger, LLC, New York, NY, (2019).
43. W. Humphrey et al., VMD: Visual molecular dynamics. J. of Mol. Graph. 14, 33-38 (1996). doi: 10.1016/0263-7855(96)00018-5
44. A. Lambiable, *et al.,* PEP-FOLD3: faster de novo structure prediction for linear peptides in solution and in complex. Nucleic Acids Res. 44, 449-54 (2016). doi: 10.1093/nar/gkw329
45. S. M. Douglas *et al.,* Rapid prototyping of 3D DNA-origami shapes with caDNAno. Nucleic Acids Res. 37, 5001-6 (2009). doi: 10.1093/nar/gkp436
46. J. Yoo, *et al.*, cadnano to PDB File Converter. https://nanohub.org/resources/cadnanocvrt. doi: 10.4231/D3SB3X05F
